# Periodic hypothalamic attractor-like dynamics during the estrus cycle

**DOI:** 10.1101/2023.05.22.541741

**Authors:** Mengyu Liu, Aditya Nair, Scott W. Linderman, David J. Anderson

## Abstract

Cyclic changes in hormonal state are well-known to regulate mating behavior during the female reproductive cycle, but whether and how these changes affect the dynamics of neural activity in the female brain is largely unknown. The ventromedial hypothalamus, ventro-lateral subdivision (VMHvl) contains a subpopulation of VMHvl^Esr1+,Npy2r-^ neurons that controls female sexual receptivity. Longitudinal single cell calcium imaging of these neurons across the estrus cycle revealed that overlapping but distinct subpopulations were active during proestrus (mating-accepting) vs. non-proestrus (rejecting) phases. Dynamical systems analysis of imaging data from proestrus females uncovered a dimension with slow ramping activity, which generated approximate line attractor-like dynamics in neural state space. During mating, the neural population vector progressed along this attractor as male mounting and intromission proceeded. Attractor-like dynamics disappeared in non-proestrus states and reappeared following re-entry into proestrus. They were also absent in ovariectomized females but were restored by hormone priming. These observations reveal that hypothalamic line attractor-like dynamics are associated with female sexual receptivity and can be reversibly regulated by sex hormones, demonstrating that attractor dynamics can be flexibly modulated by physiological state. They also suggest a potential mechanism for the neural encoding of female sexual arousal.

## Main

Fluctuating levels of sex hormones regulate an array of preparatory changes in the female body for reproduction, alter female behaviors and coordinate sexual receptivity with fertility^1–3^. Despite this well-established coordination, how hormonal states alter the computations performed by the female brain to control behavioral decisions in response to male cues is poorly understood. Estrogen Receptor 1 (Esr1)/Progesterone Receptor (PR)-expressing neurons in the ventro-lateral subdivision of the ventromedial hypothalamic nucleus (VMHvl) are known to control sexual receptivity in females^4–8^, but how they do so is not clear. Bulk calcium imaging studies have indicated that these neurons do not change their activity during the estrus cycle; rather sex steroids promote an expansion of VMHvl output fibers to AVPV^9^. More recently, however, a small subset of VMHvl^Esr1^ neurons expressing the CCK receptor (*Cckar*) has been shown to exhibit hormone-dependent changes that are correlated with mating receptivity^10^. Thus, there are conflicting data regarding whether changes in VMHvl^Esr1^ neuronal activity patterns during the estrus cycle underly cyclical changes in female sexual behavior.

To address this issue systematically, we sought to characterize neural population representations in female VMHvl during social interactions with males under different hormonal conditions. We focused on a subpopulation of Esr1^+^ neurons that are Npy2r^-^, called “α cells,” which are functionally essential for and able to promote sexual receptivity^7^. We performed longitudinal single cell imaging of VMHvl α cells in naturally cycling or ovariectomized hormone-primed females during unrestrained interactions with a sexually experienced male. By modeling VMHvl activity as a dynamical system, we identified hormonal state-dependent line attractor-like dynamics in neural state space that recur periodically during the estrus cycle, and whose presence or absence correlates with accepting vs. rejecting behavioral states, respectively.

## Tuning properties of VMHvl α neurons in females during interaction with a male

To reveal what features are encoded by female VMHvl during a mating interaction, we imaged VMHvl^Esr1+,Npy2r-^ (“α”) cells^7^ in sexually receptive females expressing GCaMP6m using a miniature head-mounted microscope^11^ (Fig. 1a). Initially, we introduced a toy, a female or a male to the tested females in their home cage (3 trials each, 1 min per trial) (Fig. 1b). We observed that distinct subsets of α cells were tuned to male or female conspecifics (Fig. 1c). Male-preferring cells were ∼4 times more abundant than female preferring cells (Fig. 1d). The average response over all neurons to a male was ∼4 times higher than the response to a female (Extended Data Fig. 1a), which is consistent with bulk calcium imaging results^7^. Principal component analysis (PCA) revealed that the first two principal components are dominated by the sex of the intruder (Extended Data Fig. 1b).

**Fig. 1.**
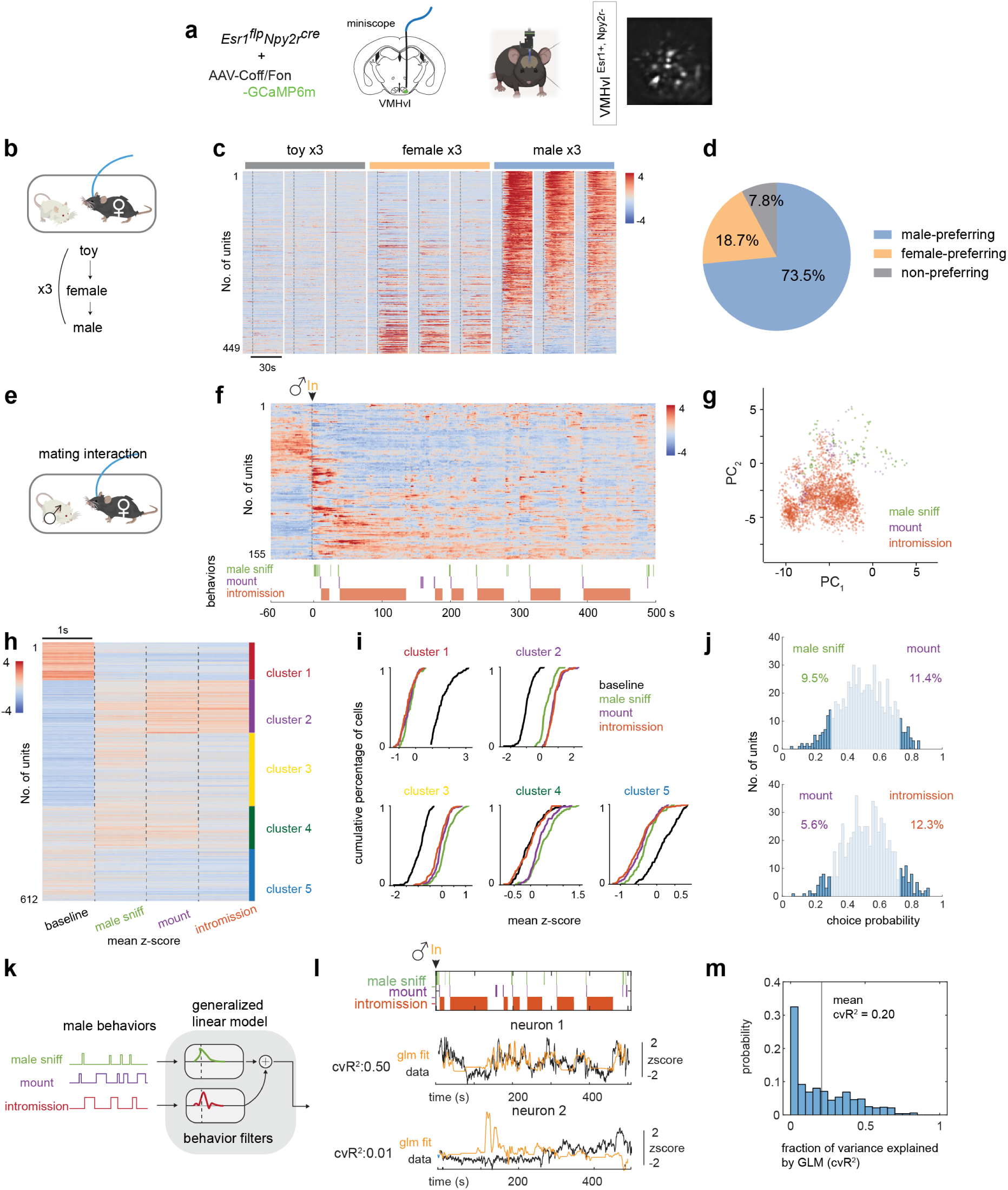
Tuning properties of VMHvl α neurons in females during interaction with a male. **a**, Left, schematic illustrating miniscope imaging of female VMHvl^Esr1+,Npy2r-^ (α) cells; Right, example imaging plane. **b**, Diagram of sex representation test. Each intruder was presented for 1 min. **c,** Concatenated average responses to toy, female, or male. N = 8 mice. Color scale indicates z-scored activity. Units were sorted by temporal correlation. **d,** Percentages of male-or female-preferring cells (calculated by Choice Probability, Methods). **e**, Diagram illustrating mating interaction test. **f**, Single cell responses during mating interactions (top) and their corresponding behaviors (bottom), from one example female. Units were sorted by temporal correlation. **g,** PCA of activity during interaction with a freely behaving male exhibiting sniffing, mounting or intromission towards female. **h,** Unsupervised clustering of female VMHvl α unit activity during mating interaction. For comparative purposes, all imaging frames (@10Hz) from each animal containing a given annotated behavior were concatenated and binned into 1 s intervals; responses of each unit were then averaged across all intervals and plotted as a single 1 s bin for each behavior. N = 6 mice. Units were sorted by cluster membership. Color scale indicates z-scored activity. **i**, Cumulative distribution of mean neuronal activity in individual clusters during indicated social behaviors; data from **h. j**, Choice Probability (CP) histograms and percentages of tuned cells. cutoff: CP>0.7 or <0.3 and >2σ. N = 6 mice. **k,** Schematic showing generalized linear model (GLM) used to predict neural activity from behavior. **l,** Example fit of selected neurons with cvR^2^ (0.50 and 0.01); behavior filters below. **m,** Distribution of cvR^2^ across all mice (N = 6 mice, mean: 0.20). Most neurons are not well-fit by the GLM. See also Extended Data Fig.1e, f.

We next performed recordings from females during free mating interactions with a male over a 10 min period (Fig. 1e, f; 64-155 units per mouse, N = 6 mice; Methods). Because females exhibit little motor behavior during mating, to investigate the correlation between VMHvl α cell activity and behavior we focused on actions performed by the male towards the female (male sniffing, mounting or intromission). We performed unsupervised clustering (Methods) of neural activity in females during male mating behaviors, and constructed raster plots of average activity during 1s intervals (Fig. 1h and Extended Data Fig. 1c). This analysis yielded distinct clusters some of which showed increased (clusters 2,3,4), while others showed decreased (cluster 1,5), activity in relation to pre-mating baseline (Fig. 1i, black lines).

Notably, activity during sniffing, mounting and intromission was not well differentiated within clusters (Fig. 1i). Consistent with this result, choice probability analysis^12^ revealed that a relatively small percentage of VMHvl α cells (∼5-12%) were “tuned” to specific male behaviors (Fig. 1j, dark blue bars), while the majority exhibited “mixed selectivity” (Fig. 1j, light blue bars), indicating relatively weak behavior-specific tuning at the population level. Consistent with this finding, PCA indicated that the activity representations of different male mating actions were not well separated from each other by the first two PCs (Fig. 1g).

To further investigate the relationship between male mating behaviors and female VMHvl neural activity, we attempted to fit Generalized Linear Models (GLMs) to the activity of each neuron as a weighted sum of sniffing, mounting and intromission behavior (Fig. 1k). The results indicated that some cells were relatively well-fit by GLMs (cross validated R^2^ (cvR^2^) ≥ ∼0.5), while others were not (cvR^2^ << ∼0.5, Fig. 1l and Extended Data Fig. 1e,f). Over all animals, only ∼20% of the variance in neural activity was explained by male mating behaviors (Fig. 1m, mean cvR^2^=0.20, N = 6 mice). Interestingly, there appeared to be a bi-modal distribution among mice in the fraction of variance explained by GLM fits in each mouse (Extended Data Fig. 1d).

## Dynamical system modeling of female VMHvl α cell activity during mating

VMHvl is well-known to exhibit sexual dimorphisms in gene expression, cell types, projections and function^6, 13^. Recently, we reported that dynamical systems modeling of male VMHvl^Esr1^ neuronal activity during aggression^12, 14^ revealed an approximate line attractor, activity along which increased as the intensity of aggressive behavior escalated from investigative sniffing to attack^15^. Because the overall variance in female VMHvl activity during mating was not well-fit by GLMs regressed against male behavior (Fig. 1m), we investigated whether this activity might be better explained by a dynamical system model. To this end, we used recurrent switching linear dynamical systems (rSLDS) analysis^15, 16^ to fit a model of α cell activity to data from each mouse (N= 6 mice). Importantly, this analysis was carried out purely on neural data, without reference to any behavioral annotation.

An rSLDS analysis yields a low-dimensional representation of neural activity, similar to principal components analysis (PCA). Unlike PCA, however, an rSLDS also segments the neural activity into discrete states with linear dynamics. An rSLDS may have purely intrinsic dynamics, or it may take sensory and behavioral covariates as input. The former procedure identified one state in isolated females lasting the duration of the period prior to male contact (state 1) and a second, long-lived neural state during mating spanning multiple male actions (male sniff, mount or intromission; Fig. 2a,b, state 2; Extended Data Fig. 2b,c; state 2 epoch duration: 742±130.5s, N = 4 mice). The fit to the data of the rSLDS models was examined using two quantitative metrics^15^: variance explained and forward simulation accuracy (FSA = 1-[normalized m.s.e. of forward simulation]). We obtained good rSLDS model fits (FSA mean=0.92 **±** 0.01) in 4/6 animals (mice 1-4) (Fig. 2a and Extended Data Fig. 2b,c,j). The variance in neural activity explained by rSLDS models in these 4 mice was high (68.75%; Extended Data Fig. 2a, 3f, “w/o beh inputs”) and was only marginally increased (∼10%) by re-fitting the rSLDS model using a male behavioral input term (Extended Data Fig. 3f,g; pink bar), indicating that the neural dynamics in these animals was largely independent of input, i.e., intrinsic.

**Fig. 2.**
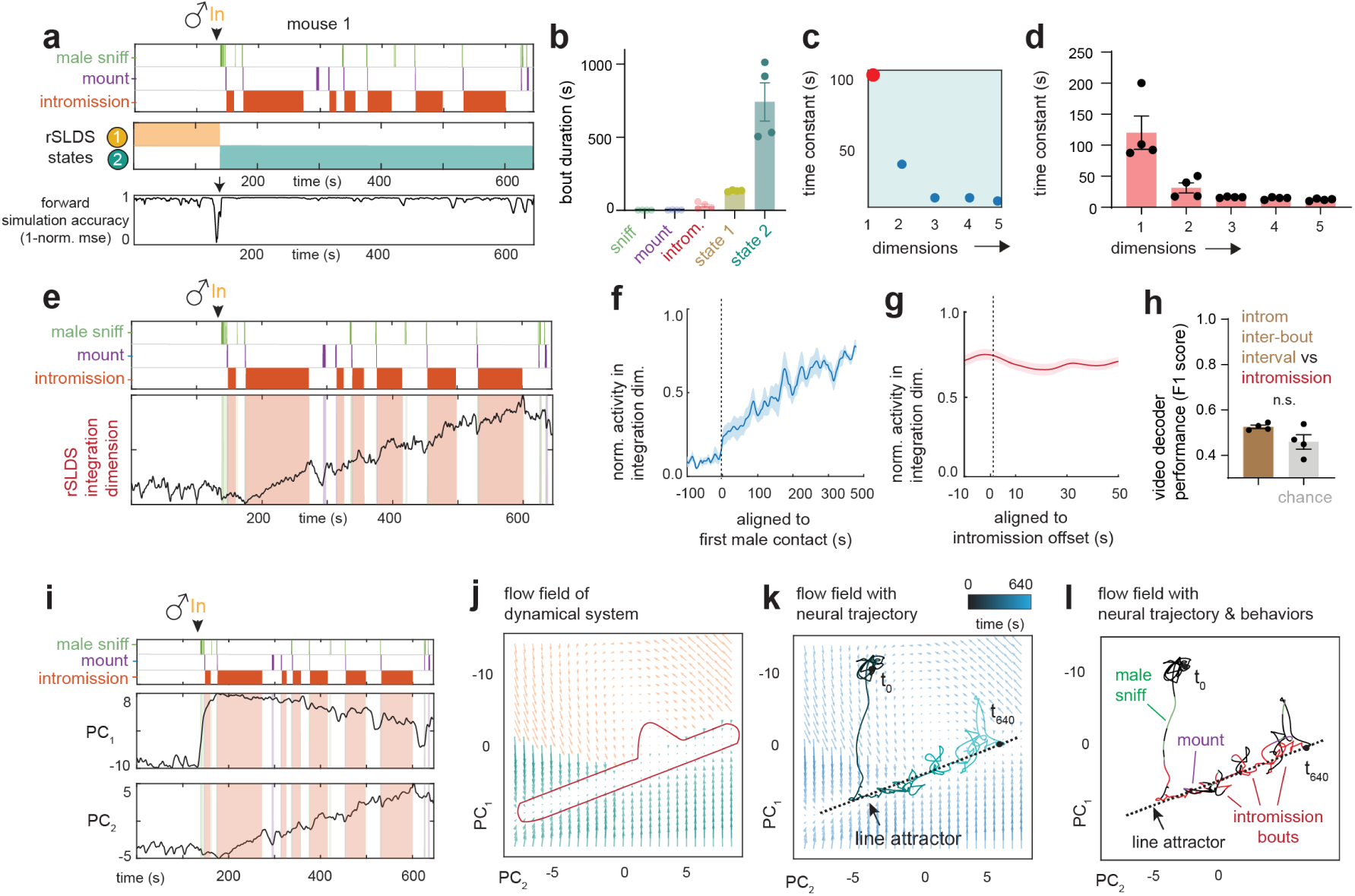
Dynamical system modeling of female VMHvl α cell activity during mating. **a**, Top: States discovered by recurrent switching linear dynamical systems (rSLDS) aligned to behaviors performed by the male in example mouse 1. Bottom: Model performance measured by forward simulation accuracy^15^ in example mouse 1. **b**, Time scale of individual bouts of behaviors versus time scales of rSLDS states (N = 4 mice). **c,** Time constants of state 2 (cf. **a**) reveal a single dimension with a large time constant in mouse 1. **d,** Distribution of time constants across animals well-fit by the rSLDS model (N = 4 mice). Time constants sorted by magnitude in each animal. **e**, Dynamics of integration dimension in mouse 1 reveals a ramping dimension that correlates with escalating sexual arousal. **f**, Behavior triggered average of the normalized activity of the integration dimension aligned to the onset of male contact. (N = 4 mice). **g,** Behavior triggered average of the normalized activity of the integration dimension aligned to the offset of intromission. (N = 4 mice). **h**, Videoframe behavioral decoder performance trained on neural data from intromission inter-bout interval vs. intromission. (N = 4 mice). **i,** Low dimensional principal components of VMHvl α cell dynamical system. **j,** Flow field of VMHvl α dynamical system colored by rSLDS states. **k,** Flow field of VMHvl α dynamical system showing neural trajectories in state space. **l,** Neural state space of VMHvl α dynamical system highlighting regions where fixed points are present (dash line).

**Fig. 3.**
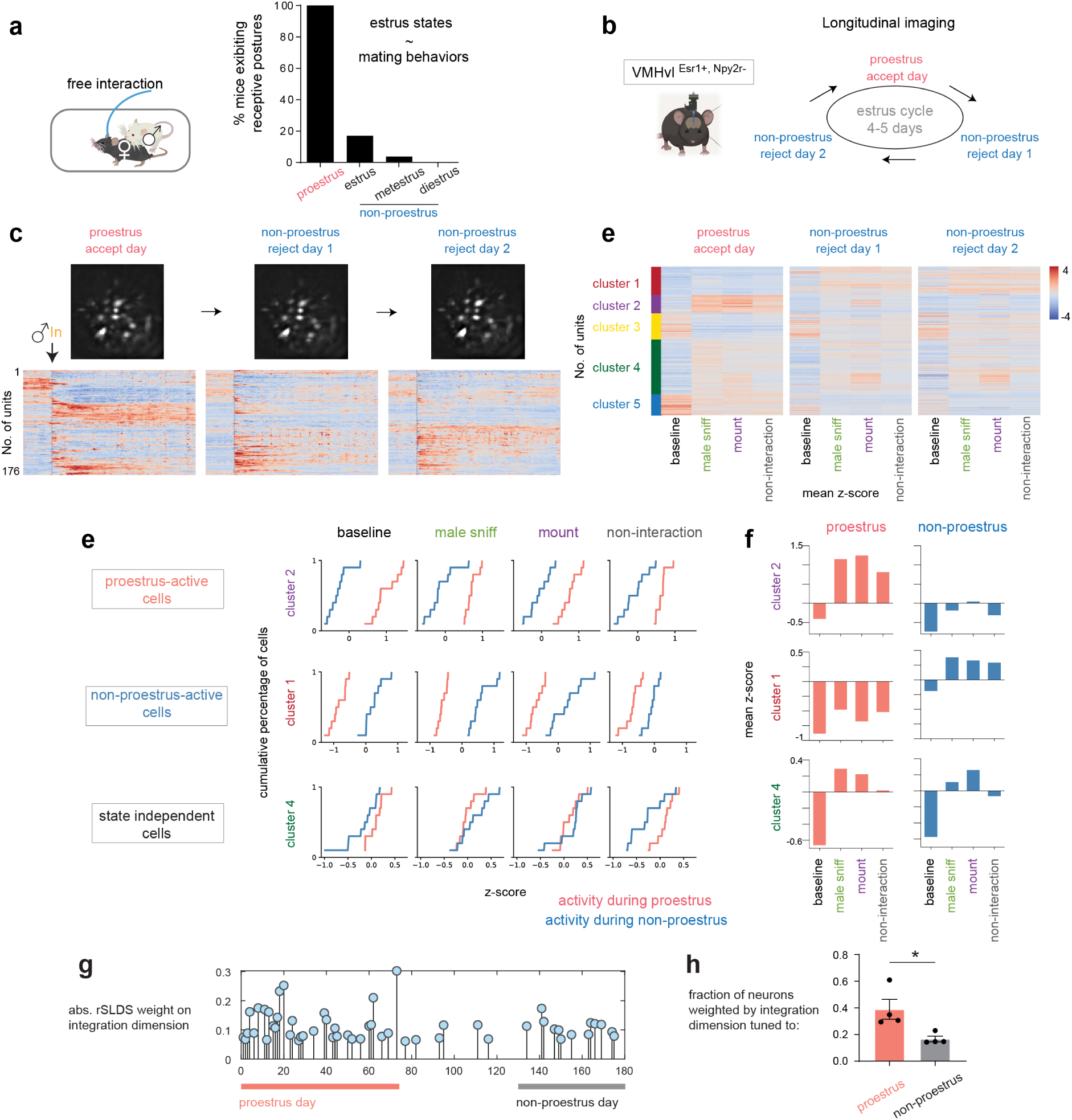
Population activity in VMHvl α cells changes across the estrus cycle. **a**, Correlation between female estrus states and mating behaviors. **b,** Illustration for estrus states of naturally cycling females and longitudinal imaging strategy. **c,** Example longitudinal imaging planes and traces from one female. Units were sorted by temporal correlation. Color scale indicates z-scored activity. **d,** Unsupervised clustering of neuronal activity during mating interactions across estrus states. Combined data from N = 6 mice. Units z-scored across days. **e,** Cumulative distribution of mean neuronal activity in individual clusters during different social behaviors. Non-interaction indicated the time when the bodies of the two mice were physically separated. Colors indicate proestrus vs. pooled non-proestrus states. **f,** Mean activity of selected clusters during indicated social behaviors. **g,** Absolute weights contribution of individual neurons to the integration dimension in mouse 1 sorted by neurons’ tuning towards the proestrus or non-proestrus phase. **h,** Fraction of neurons weighted by the integration dimension with tuning towards proestrus vs non-proestrus day. N = 4 mice.

In 2 of the 6 animals imaged (mice 5 and 6), an rSLDS with purely intrinsic dynamics yielded poor fits (3.8-5.8% variance explained, Extended Data Fig. 2a,n). Strikingly, the GLM fits for these 2 mice (i.e., variance explained by male contact behaviors) was higher than for the 4 mice well-fit by rSLDS (Extended Data Fig. 3a-c; mean cvR^2^= 0.35 vs. 0.15, respectively). Accordingly, among all 6 mice, there was a strong inverse correlation (R^2^=0.93) between variance explained by GLM fits vs. rSLDS fits (Extended Data Fig. 3e). Furthermore, in the 2 mice not well-fit by rSLDS, including a male behavioral input term in the model dramatically increased the variance explained (Extended Data Fig. 3f,g). Thus, in these two mice variance explained by dynamics is primarily input-driven rather than intrinsic. Potential reasons for this biological variation will be discussed below (Extended Data Fig. 9).

In those mice with good rSLDS model fits (N = 4 mice; n=137±48 neurons/mouse; Extended Data Fig. 3f), examination of the time constants (Eigenvalues) of the dynamics matrix fit to the neural data in state 2 revealed that one dimension had a very long time constant in relation to the other dimensions (Fig. 2c, red dot). The average value of this time constant across animals was 120.3±26.72s (Fig. 2d, dimension 1; N = 4 mice). We examined the temporal correlation between the average activity of the neurons weighted by this slow dimension and male mating behaviors. Activity began to increase at the onset of male contact-mediated behaviors (intromission, mounting or sniffing, depending on the mouse) and continued to ramp up throughout multiple male-contact bouts (Fig. 2e,f, Extended Data Fig. 2d,e,k, N = 4 mice). Dimensions with long time constants can serve as integrators^17^, inputs can push the neural state along this dimension and the state can persist long after. The observed ramping activity through male-contact bouts is reminiscent of such an integrator, so we operationally refer to this dimension as the “integration dimension”^15^.

Male mice perform intromission in bouts separated by inter-bout intervals (IBIs; Fig. 2a, red vs. white rasters, respectively). Notably, the average value of the rSLDS integration dimension during these IBIs was relatively high, and similar to that during the preceding intromission bout (Fig. 2g). Accordingly, decoders trained on the scalar value of the integration dimension to distinguish videoframes recorded during intromission bouts vs. IBIs performed no better than chance (Fig. 2h). Because the male is physically separated from the female during IBIs, these data suggest that activity along the integration dimension does not simply encode periodic male contact-dependent sensory input, but rather represents a signal that is gradually accumulated or integrated over time (Fig. 2e and Extended Data Fig. 2d,e,k).

Inspection of the neurons that contributed the highest weights to the integration dimension revealed that a relatively small percentage of these cells (8%∼14%) were “tuned” to specific male behaviors (Extended Data Fig. 4a, dark blue bars), while the majority exhibited “mixed selectivity” (Extended Data Fig. 4a, light blue bars), similar to the population as a whole (Fig. 1j). Analysis of individual traces of such neurons indicated that some cells with ramping-like activity were observed (Extended Data Fig. 4d-f, cells n_38_, n_41_, n_25_) and that different cells peaked at different times during the male interaction (Extended Data Fig. 4b,c, orange arrowheads). This suggests that the progressive ramping up of activity in the integration dimension (Extended Data Fig. 4b, *upper*) may be an emergent property of the population, at least in part. In addition, calculation of the halfwidth of each neuron’s autocorrelation function (ACHW) (which value is an approximation of the neuron’s time constant^18, 19^) identified some cells exhibiting apparent persistent activity across inter-intromission bout intervals (Extended Data Fig. 4g-i). Notably, the ACHW of the same cells significantly decreased when the male was confined in a perforated pencil cup during free mating interaction (Extended Data Fig. 4j-m). This result suggests that the persistent activity observed across sporadic intromission bouts and their IBIs cannot be explained simply as cells modulated by male odors.

We sought next to visualize the features of the dynamical system in a low-dimensional state space. The log_2_ of the ratio of the time constant of the integration dimension to that of the next-slowest dimension (Fig. 2d, dimension 2), a parameter we call the “line attractor score”^15^, was significantly >0 in all 4 mice (mean value: 2 **±** 0.23, p **<** 0.001). The high value of this parameter as well as the absolute magnitude of the time constant suggested the existence of line attractor-like dynamics in neural state space^15^. To visualize the rSLDS states in 2D, we extracted the top two rSLDS dimensions that accounted for the most variance in the neural activity (analogously to PCA). The second ‘principal component’ (PC2) of the rSLDS exhibited ramping activity (Fig. 2i; cf. Fig. 2e). Visualizing the flow field of the dynamical system revealed a stable region of state space resembling an approximate line attractor (Fig. 2j,k and Extended Data Fig. 2f,h,l). Observing neural trajectories in this state space after annotation of corresponding male behavior, we found that the neural activity vector entered the line attractor following initial close contact with the male (Fig. 2l and Extended Data Fig. 2g,i,m), and progressed along it during successive male mating bouts. This progression reflects the ramping seen in the integration dimension that contributes to this attractor (Fig. 2e and Extended Data Fig. 2d,e,k). These data suggest that VMHvl α cell activity displays approximate line attractor-like or leaky integrator-like dynamics during mating.

**Fig.4.**
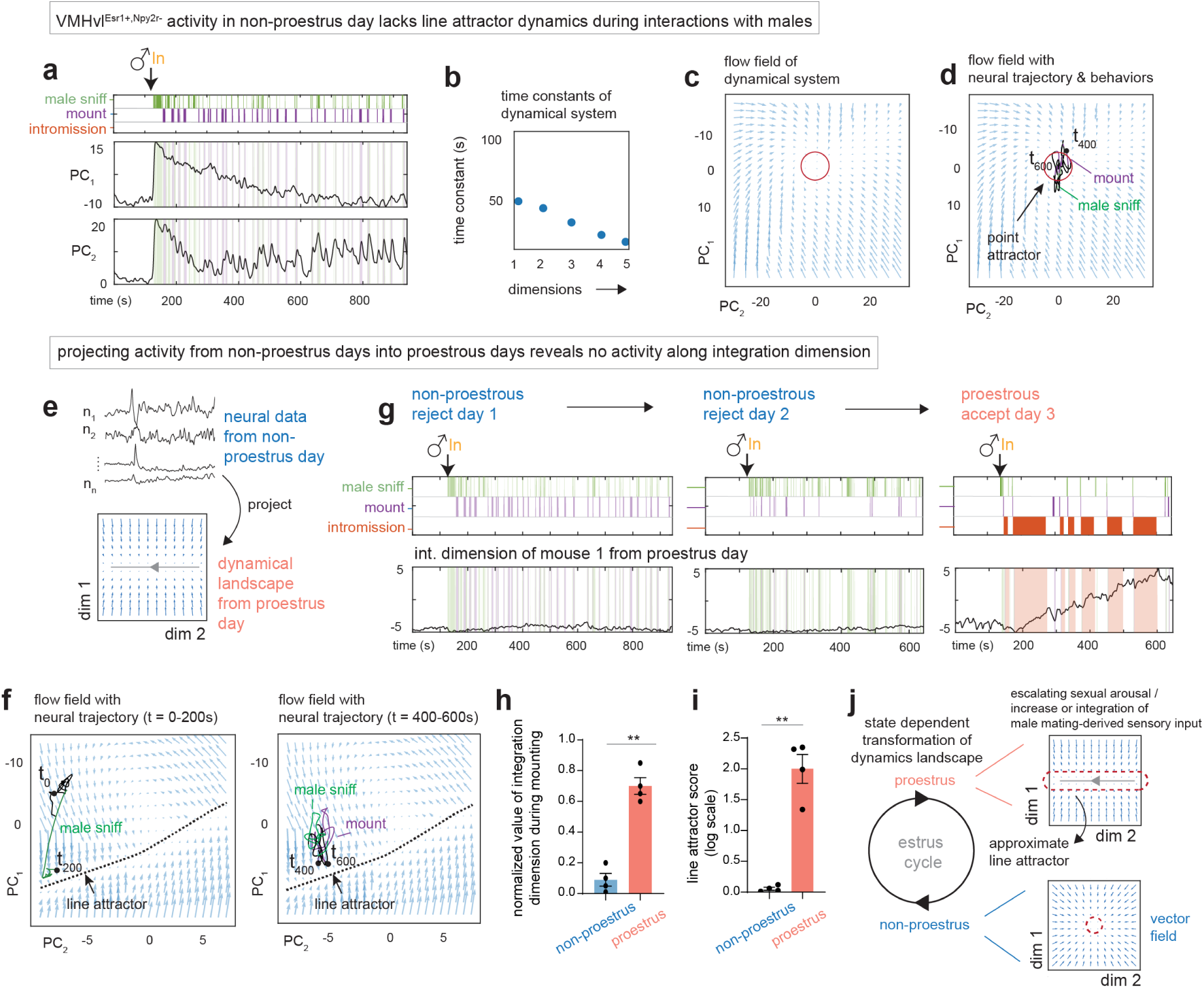
Neural dynamics in female VMHvl is modulated by the estrus cycle. **a**, Low dimensional principal components of VMHvl α rSLDS fit model in non-proestrus day of mouse 1. Principal components show fast time locked dynamics and lack ramping and persistence. **b,** Time constants of VMHvl α dynamical system in non-proestrus day for mouse 1. **c,** Flow field of VMHvl α dynamical system in non-proestrus day. **d,** Same as **c**, showing neural trajectories in state space colored by time and colored by behaviors (right). **e,** Schematic illustrating projection of neural activity from non-proestrus day into fit dynamical system from proestrus day. **f,** Flow field of VMHvl α dynamical system in proestrus day with neural trajectories projected from non-proestrus day for t = 0 to t = 200s (left) and t = 200s to t = 400s (right). **g,** Dynamics of integration dimension in VMHvl mouse 1 discovered during proestrus day compared to activity of the same dimension on non-proestrus days. **h,** Quantification of normalized value of integration dimension during male-mounting in non-proestrus and proestrus days (N = 4 mice). **I,** Line attractor score for dynamical systems fit during proestrus and non-proestrus days (N = 4 mice). **j,** Schematic highlighting state dependent transformation of neural dynamics during the estrus cycle.

## Population activity in VMHvl α cells changes across the estrus cycle

Females adjust their mating behaviors (decide whether to accept or reject a male) according to their hormonal state. We first confirmed the correlation between estrus state in naturally cycling females (estimated by vaginal smears) and mating behaviors (acceptance vs. rejection). Females were sexually accepting in the proestrus state, while rejecting in metestrus, diestrus and estrus (non-proestrus states; Fig. 3a; rejection is not passive but involves active avoidance actions such as kicking). We next sought to identify neural correlates of this change in behavior. Consistent with previous studies^7, 9^, bulk calcium recording revealed no estrus cycle phase-related changes in VMHvl α cell population activity during interactions with males (Extended Data Fig. 5a). However, complementary changes in VMHvl neuron population activity between mating-accepting vs. rejecting states could cancel each other out, leading to no net change in average activity. To examine this possibility, we performed longitudinal single-neuron-resolution imaging in VMHvl α cells in females during mating interactions with an unrestrained sexually experienced male. We compared neural activity in each animal across 1 proestrus/sexually accepting day and 2 non-proestrus/rejecting days (Fig. 3b,c).

Single cell activity during male encounters was negatively correlated between proestrus and non-proestrus days, while activity compared between two non-proestrus days was positively correlated (Extended Data Fig. 5b), indicating a change in population activity patterns. (Note that this change was detectable even though we omitted from the analysis activity recorded during intromission, which males do not perform towards non-proestrus females.) Next, we performed unsupervised clustering of VMHvl α cell activity concatenated by male mating behaviors^20^ (Fig. 3d). Some clusters displayed hormonal state-dependent activity: cluster 2 and 5 displayed significantly higher activity in proestrus, while clusters 1 and 3 were higher in non-proestrus states (Fig. 3e,f and Extended Data Fig. 5d). By contrast, the activity of cluster 4 showed no significant difference across hormonal states, responding to male contact regardless of hormonal states.

To achieve better control over hormonal state, we repeated the longitudinal imaging experiments in ovariectomized (OVX) females before vs. after hormone priming (oil vs. E+P in oil; OVX/EP; Extended Data Fig. 6a). Unsupervised clustering of VMHvl α cells (Extended Data Fig. 6b) yielded similar results to those observed in naturally cycling females: we observed clusters with greater activity in the OVX/EP (accepting; proestrus equivalent) state, the OVX/oil (rejecting; non-proestrus equivalent) state and those with equivalent activity in both states (Extended Data Fig. 6c,d). Together, these results indicate that the female VMHvl α cells active during interactions with males comprise overlapping but distinct subpopulations with different activity patterns at different phases of the estrus cycle.

These observations raised the question of whether the neurons that place high weight on the integration dimension of the rSLDS fit using proestrus data (cf. Fig. 2e) were also active during non-proestrus phases. To answer this question, we calculated the weights of each unit on the integration dimension during proestrus (Fig. 3g) and sorted them according to whether each unit was more active during proestrus or non-proestrus states using choice probability to classify neurons into either category. This analysis revealed that 43% of highly weighted neurons did not exhibit preferential activation during a particular hormonal state. Of the remaining highly weighted neurons, a large fraction (40%) was preferentially tuned to the proestrus state (based on choice probability), while a little less than half as many (17%) were tuned to non-proestrus states (Fig. 3h). Thus, the integration dimension of the rSLDS fit model is weighted by a similar proportion of neurons that are more active when females are sexually receptive and those that are active throughout the estrus cycle.

## Neural dynamics modulated by the estrus cycle

We investigated next whether approximate line attractor dynamics were exhibited by VMHvl α cells during non-proestrus phases, by fitting an rSLDS model to data recorded on those days. Remarkably, in contrast to the proestrus model the latent factors discovered by rSLDS did not include a single dimension with a time constant substantially larger than that of the next slowest dimension (Fig. 4b), as reflected in the low line attractor score of the fit model (Fig. 4i). Instead, we observed relatively fast dynamics time-locked to occurrences of male sniffing and mounting (PC2 in Fig. 4a). Consistent with this finding, examining the underlying flow-field of the rSLDS dynamical system fit to non-proestrus data failed to reveal evidence of a line attractor. Rather the neural state space contained a single point attractor, reflecting stable baseline activity prior to interaction with the male (Fig. 4c). During male sniffing and mounting, activity in the female brain exited the attractor briefly but quickly returned to the same fixed point (Fig. 4d, Extended Data Fig. 7a,b), reflecting the absence of persistent activity or ramping during non-proestrus.

To visualize better the neural dynamics observed during non-proestrus phases in the context of the flow field derived from the rSLDS model in proestrus, we projected neural activity from non-proestrus days into the dynamical system model fit using proestrus day data (i.e., assigned weights to the neurons active during non-proestrus days using the dynamics matrix fit to the proestrus data; Fig. 4e). In this visualization, activity during non-proestrus phases remained at one end of the line attractor throughout the interaction with males and did not show any progression along it even as attempted male mounting occurred (Fig. 4f). Correspondingly, in the PC space derived from proestrus data, the projected non-proestrus day data did not show ramping along the integration dimension (Fig. 4g, Extended Data Fig. 7d,e) even though it contains numerous active cells that weight the integration dimension of the proestrus model (Fig. 3h). Notably, although male mounting occurred on both proestrus and non-proestrus days (Fig. 4g, purple ticks), the average value of the integration dimension during this behavior was significantly lower on non-proestrus days (Fig. 4h). Thus, the failure of rSLDS to discover an integration dimension in the non-proestrus data does not simply reflect the lack of a proestrus day-specific male behavior (intromission), but instead (or in addition) reflects a difference in the intrinsic dynamics of neurons which are active during mounting on proestrus vs. non-proestrus days.

To directly test whether the change in population dynamics observed across the natural estrus cycle was due to changes in hormone levels, we repeated our dynamical systems analysis in the same females after ovariectomy (OVX) with or without hormone priming (OVX/oil vs, OVX/EP; Extended Data Fig. 8a-d). This revealed similar dynamical landscapes when rSLDS models were fit to data recorded in proestrus during natural cycling or to data from hormone-primed OVX mice (Extended Data Fig. 8a-h). Crucially, the integration dimension discovered by fitting an rSLDS model to data from the hormone primed day exhibited little activity in OVX/oil females (Extended Data Fig. 8j), directly demonstrating the hormone dependence of this latent factor.

As mentioned earlier, of the 6 females recorded during natural cycling, two exhibited a poor rSLDS model fit during proestrus (Extended Data Fig. 2n, forward simulation accuracy). In one of these animals (mouse 5; Extended Data Fig. 9a), we performed OVX and hormone priming after completing the recordings during natural cycling (the second animal was sacrificed before OVX could be performed). Recordings from this mouse during interactions with a male after OVX/hormone-priming yielded a much better rSLDS model fit than was obtained during natural proestrus (Extended Data Fig. 9a,b, forward simulation accuracy). This fit model exhibited approximate line attractor dynamics (Extended Data Fig. 9c-e). Modulation in the integration dimension was not observed after OVX but before hormone-priming (Extended Data Fig. 9f). These results suggest that the poor fit of the rSLDS model on data recorded from this mouse during natural proestrus might be due to lower hormone levels in this mouse (which cannot be directly measured in naturally cycling females), than those in mice initially well-fit by rSLDS. This conclusion is supported by the observation that even in the latter mice, the fit of the rSLDS model to OVX/EP data is better than the fit to the data recorded from the same animal on a proestrus day (Extended Data Fig. 8a,b, forward simulation accuracy).

Line attractor dynamics in other neural systems reflect persistent neuronal activity, which creates the stability characteristic of attractors^21^. To measure the degree of persistence in the recorded neurons, we calculated the halfwidth of each neuron’s autocorrelation function^12^. We then compared these values for cells that were active during both proestrus and non-proestrus phases, and which contributed to the integration dimension during proestrus (Extended Data Fig. 10). Such neurons exhibited a significantly larger autocorrelation half-width (ACHW) during proestrus, relative to that during non-proestrus days (Extended Data Fig. 10b; distribution mean for ACHW in non-proestrus state = 16.1 ± 0.8s; distribution mean for ACHW in proestrus state = 25.2 ± 1.5s, p<0.001). This difference in ACHW was observed regardless of the order in which proestrus and non-proestrus days occurred in different mice (Extended Data Fig. 10c,f). Neurons that did not contribute to the proestrus integration dimension (such as non-proestrus-preferring neurons) did not exhibit a change in ACHW across the estrus cycle (Extended Data Fig. 10d,g).

Finally, we compared the ACHW of each neuron between proestrus vs. non-proestrus states. A scatterplot of these data revealed a subpopulation of line attractor neurons whose ACHW increased during proestrus relative to their non-proestrus values (Extended Data Fig. 10e,h, red datapoints). Similar results were obtained for the same animals after OVX with or without hormone priming: the neurons that contribute to the line attractor exhibited an increased ACHW on the hormone primed day, relative to the non-primed days (Extended Data Fig. 10i-n).

Taken together, these data reveal a reversible hormone dependent change in neural dynamics within the α cell population in female VMHvl across the estrus cycle (Fig. 4j). During proestrus, the dynamical landscape defined by population neural activity contains an approximate line attractor or leaky integrator. However, this putative line attractor is absent during non-proestrus phases.

## Discussion

In many species, hormones fluctuate across the estrus cycle and control changing female sexual behavior. However, whether and how these hormone-dependent behavioral changes are controlled at the level of neural population activity is largely unknown. This knowledge gap in part reflects the traditional “siloing” of the behavioral and systems neuroscience communities. The data presented here bridge this gap by bringing single-neuron imaging and dynamical systems analysis to bear on the problem of hormone-dependent behavioral control.

Esr1^+^ neurons in VMHvl have long been known to control cyclical hormone-dependent female sexual behavior in rodents^4, 5^, but how they do so is not yet clear^6–10^. Our results have revealed an approximate line attractor that emerges from population activity in a genetically defined subset of VMHvl^Esr1^ neurons during proestrus, and which appears to encode a continuous, escalating state variable in females during mating. We note that the fit rSLDS model that yielded an approximate line attractor is agonistic to the underlying cellular mechanisms, and does not necessarily reflect the recurrent connectivity typically invoked to explain attractor dynamics in other systems^21^. More data will be required to determine whether these dynamics reflect a true line attractor^22, 23^, “leaky” integration^24^ or some other type of dynamical change. Whatever the case, our data reveal that neural activity dynamics in female VMHvl varies periodically during the estrus cycle, in a hormone-dependent manner.

Our observations are surprising, because attractor dynamics have generally been considered as stable coding properties of neural circuits^21^. We identify, to our knowledge, the first case of an approximate line attractor in neural state space that appears and disappears with a change in hormonal state. Although we have recently discovered an approximate line attractor in male VMHvl^Esr1+^ cells that appears to encode an escalating internal aggressive state^15^, there is no evidence for its state-dependence. Male aggression, unlike female aggression or mating receptivity, is constitutive rather than periodic.

Unlike the case in males, where progression along the approximate line attractor correlated with the escalation of aggressive behavior^15^ the behavioral or state feature represented by progression along the approximate line attractor in receptive females during mating is not yet clear. In part this difference may reflect the fact that in males, neural activity and behavior were correlated in the same individual animals, whereas here neural activity in females was correlated with male behavior. The ramping activity in the integration dimension could reflect a gradual increase in the intensity or frequency of male contact-mediated sensory input; the integration of such input over time; or increasing sexual arousal. Several lines of evidence support the last hypothesis. First, optogenetic activation of VMHvl α cells promoted female mating receptivity (even during diestrus), while silencing these neurons inhibited receptivity in proestrus^7^. Second, in proestrus females activity along the integration dimension remained elevated from intromission bouts into the adjacent IBIs, when the animals were physically separated from each other (Fig. 2e-h). This persistence cannot be explained by the continuous presence of male chemosensory cues during mating, since it was not observed in females exposed to males in a perforated pencil cup (Extended Data Fig. 4j-m). Finally, ramping was observed in rSLDS fits to proestrus data with or without an input term representing male behavior (Extended Data Fig. 3h-k), suggesting it reflects dynamics intrinsic to the female. A role for a line attractor in representing increasing sexual arousal fits the intuition that arousal is a continuous variable that escalates during mating. By contrast, sexual rejection behavior during non-receptive phases is sporadic and punctuated; therefore slow dynamics are not necessary.

The nature of the different α cell subpopulations that underly cycling attractor dynamics remains to be determined. Our previous work has shown that transcriptionally distinct subsets of VMHvl^Esr1^ neurons, called α and β cells, control female sexual receptivity and maternal aggression, respectively^7^. Here we show that the α cell population exhibits further heterogeneity at the physiological level, comprising hormonal state-dependent and-independent subpopulations. Whether these subpopulations are transcriptomically distinct is not yet clear^10, 25^ and will require further studies. Recently, it was reported that a subset of the α cell population expressing Cckar (VMHvl^Cckar^ neurons) displayed reproductive state dependent changes in spontaneous activity and responsivity to male cues as measured by fiber photometry, with the highest levels observed during estrus. Furthermore, in vitro experiments revealed a state-dependent alternation of excitability and synaptic inputs^10^. Whether the VMHvl^Cckar^ cells and their state-dependent changes contribute to the reversible line attractor like dynamics observed here requires further investigations.

Our results demonstrate that neural population dynamics can be reversibly sculpted by hormonal influences on the brain. Because the cellular, molecular and connectional features of VMHvl are well-described^6–8, 13, 26, 27^, this system may provide an opportunity to understand how hormones, genes, cell-types and local circuitry generate emergent neural population dynamics that underlie innate social behaviors.

---

**Extended Data Fig.1.**
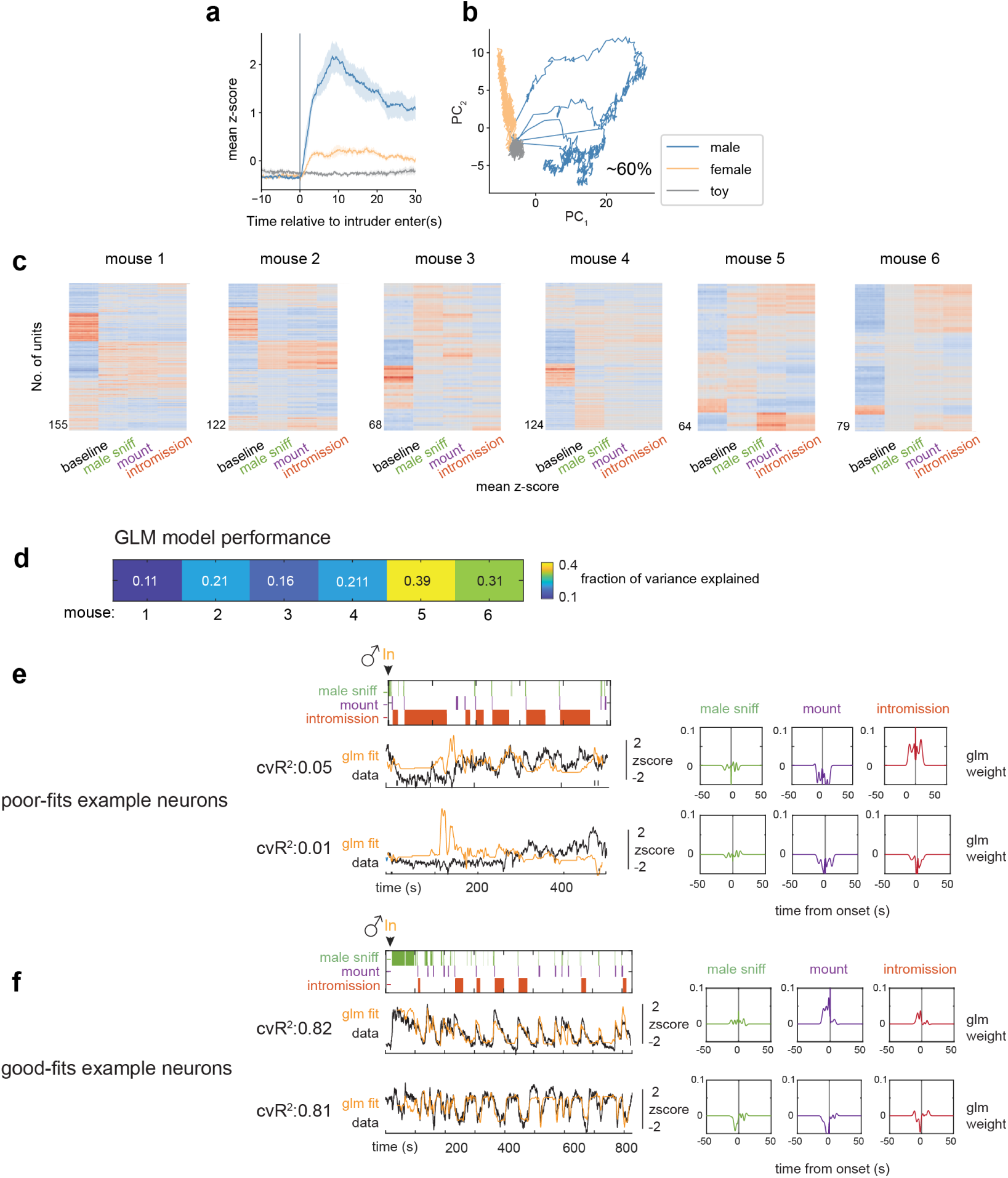
Additional information for Fig. 1. **a**, Mean responses of female VMHvl^Esr1^ α cells to male, female and toy. N=8 mice. **b,** PCA of neuronal responses to male, female and toy from one example female. **c,** Unsupervised clustering of neuronal activity from individual mice. Columns represents activity averaged across 1s windows during each indicated behavior. **d,** GLM model performance (mean fraction of variance explained across all neurons) in each mouse. **e,f,** Example generalized linear model fits and behavior filters for poorly fit neurons **e**, from mouse 1 and well fit neurons **f**, from mouse 5.

**Extended Data Fig.2.**
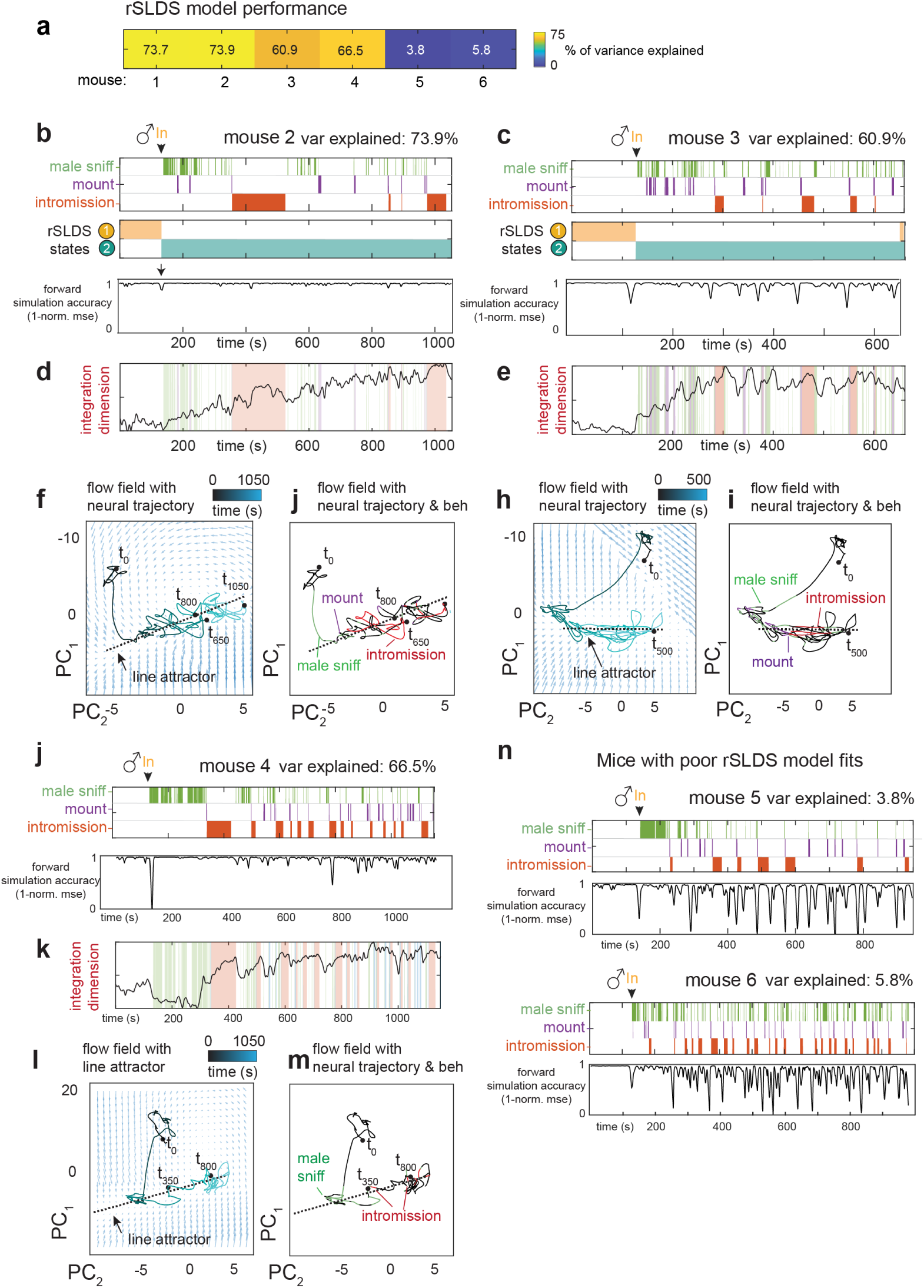
rSLDS model fitting for all mice, additional information for Fig.2. **a**, rSLDS model performance (mean fraction of variance explained across all dimensions in given mouse) in each mouse. **b,** States discovered by recurrent switching linear dynamical systems (rSLDS) aligned to behaviors performed by the male intruder towards female mouse 2 (Top). rSLDS model fit (forward simulation accuracy, bottom). **c,** Same as **b** for mouse 3. **d,** Dynamics of integration dimension in mouse 2. **e,** Same as **d** for mouse 3. **f,** Flow field of VMHvl α dynamical system for mouse 2 showing neural trajectories in state space, annotated by time from male encounter (t_0_). **g,** Neural state space of VMHvl α dynamical system for mouse 2 highlighting male behaviors and region containing approximate line attractor. **h,** Same as **f** for mouse 3. **i,** Same as **g** for mouse 3. **j,** rSLDS model fit for mouse 4. **k,** Dynamics of integration dimension in mouse 4. **l,** Same as **f** for mouse 4. **m,** Same as **g** for mouse 4. **n,** rSLDS model fit for mouse 5 and 6 highlighting a poor model fit in these animals.

**Extended Data Fig.3.**
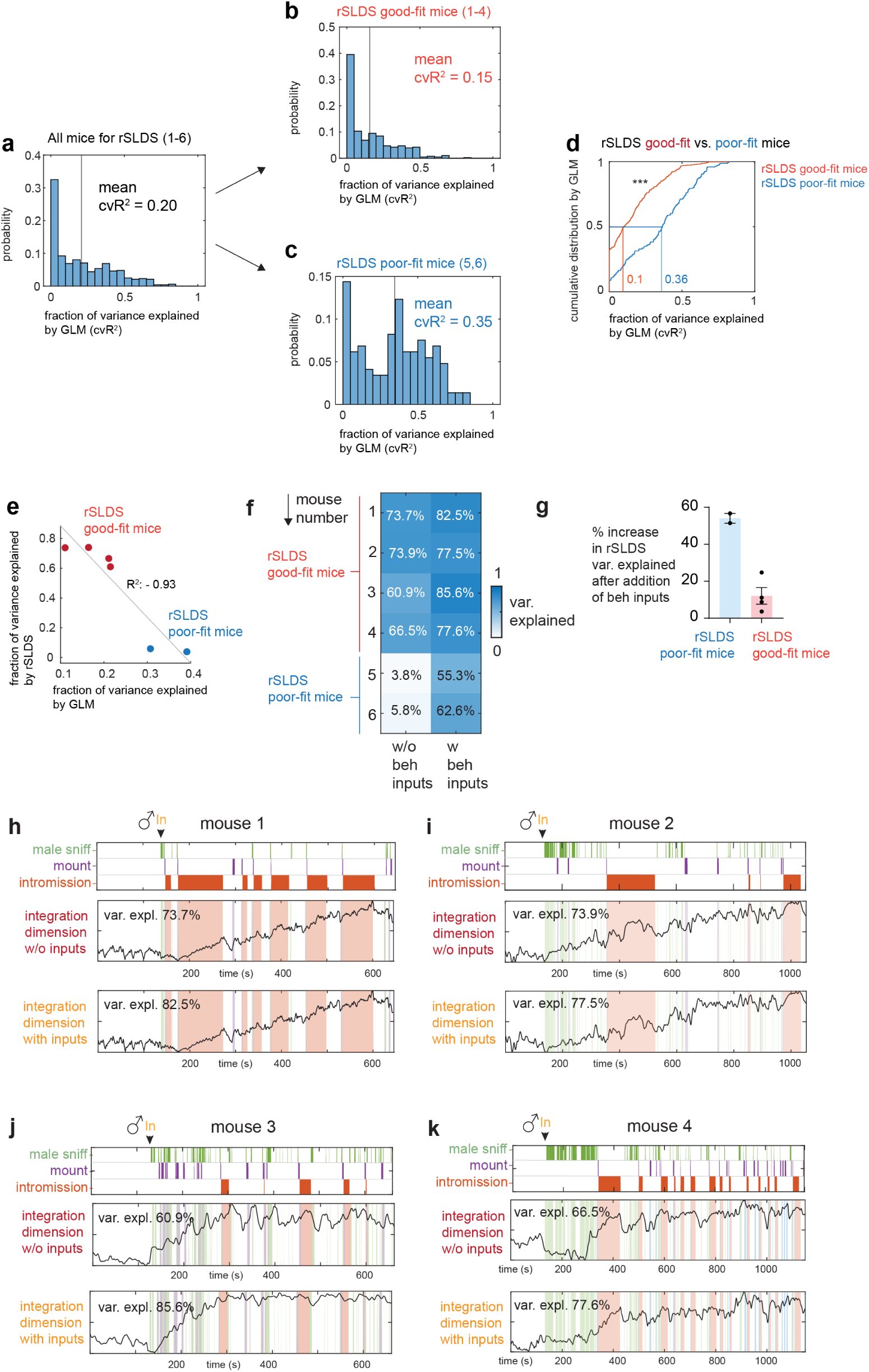
GLM fits for rSLDS good-fit and poor-fit mice, additional information for Fig.2. **a**, Variance explained by GLM trained to predict neural activity from behavior for all mice used for rSLDS (distribution mean = 0.20, N = 6 mice). **b,** Variance explained by GLM trained to predict neural activity from behavior for mice with good rSLDS fits (distribution mean = 0.15, N = 4 mice, mouse 1-4). **c,** Variance explained by GLM trained to predict neural activity from behavior for mice with poor rSLDS fits (distribution mean = 0.35, N = 2 mice, mouse 5, 6). **d,** Cumulative distribution of GLM model fit (cross validation R^2^) for rSLDS good-fit mice (median: 0.1) vs rSLDS poor-fit mice (median: 0.36), ***p<0.0001. **e,** Correlation between variance explained by GLM and rSLDS. **f,** Variance explained by rSLDS fitting with or without behavior inputs. **g,** Percentage of increase in rSLDS variance explained after adding behavior inputs. **h,** rSLDS integration dimension in models fit with and without male behavior inputs in mouse 1. The rSLDS model performance as variance explained is also shown for each model. Percentage of increase in rSLDS variance explained after adding behavior inputs. **i,** Same as **h,** for mouse 2. **j,** Same as **h** for mouse 3. **k,** Same as **h** for mouse 4.

**Extended Data Fig.4.**
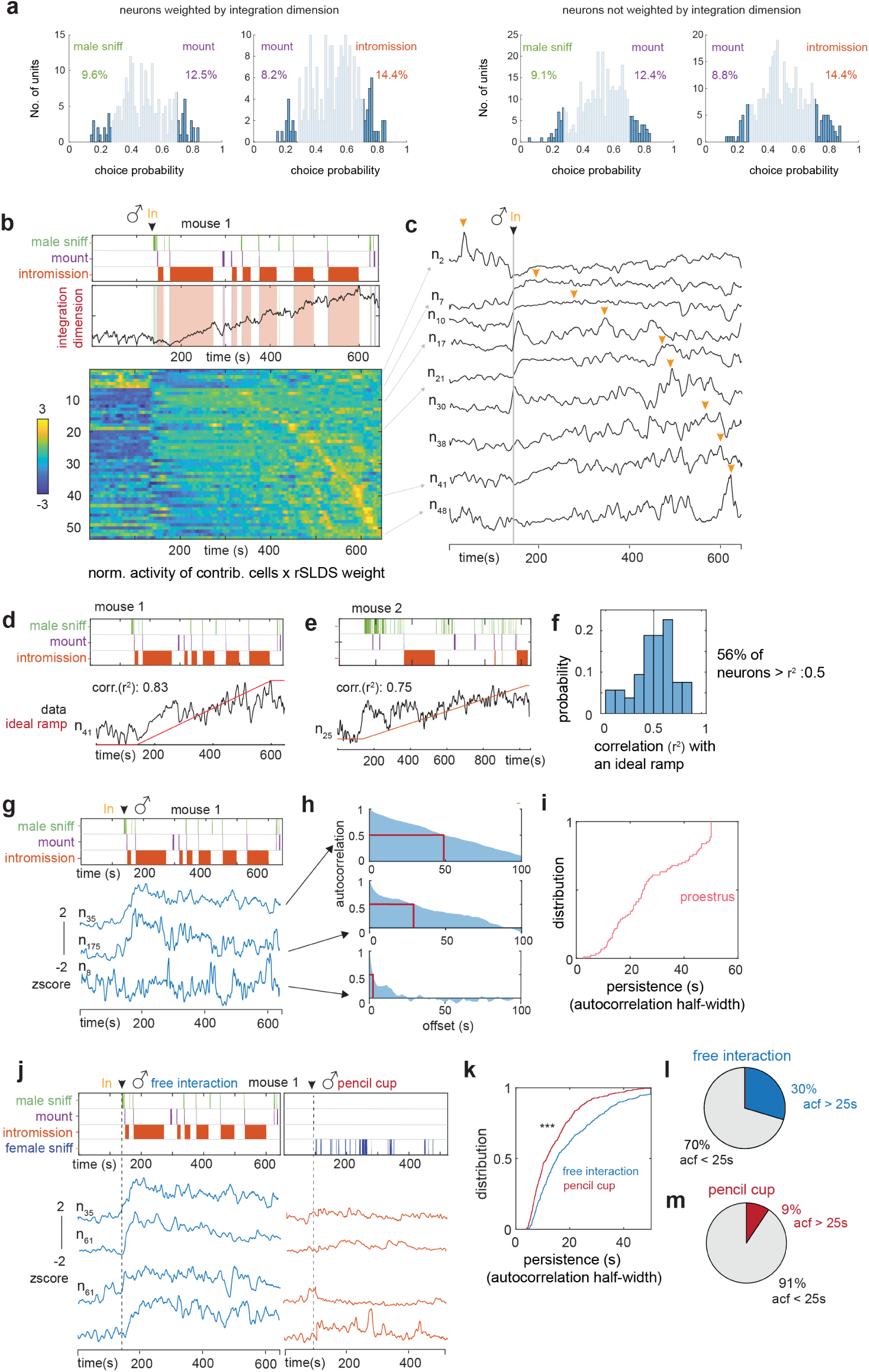
Properties of line attractor cells, additional information for Fig.2. **a**, Choice Probability (CP) histograms and number of tuned cells from highly weighted (left) cells or not highly weighted cells (right) to integration dimension. cutoff: CP>0.7 or <0.3 and >2σ. N=4 mice **b, *Upper***, relationship of male behavior to weighted average of all units contributing to integration dimension as a function of time. Data from 1 example mouse. ***Lower***, normalized activity (z-score) of individual units times rSLDS weight for each unit exhibiting a significant weight in the integration dimension of VMHvl mouse 1, sorted by time to peak. **c,** Traces of example units from **b**, *lower*. Yellow arrow indicates peak of activity for each unit. **d,** Correlation of example units activity with an ideal ramp in mouse 1. **e,** Same as D for example unit in mouse 2. **f,** Distribution of correlation of individual neuron activity with ideal ramp. **g,** Example VMHvl neurons in proestrus day showing a range of persistent activity (z-scored ΔF/F). **h,** Auto-correlation half-width (ACHW) as a measure of persistent activity, for example units shown in **g. i,** Cumulative distribution of ACHW for units with significant weights on the integration dimension during proestrus day (N = 4 mice). **j,** Dynamics of persistently active neurons identified during proestrus day during pencil-cup assay. **k,** Cumulative distribution of ACHW for same neurons during free interaction vs pencil cup assay ***p<0.001. **l,** Pie chart indicating fraction of neurons with ACHW > 25s in free interaction. **m,** Same as **l** for data obtained during pencil cup assay.

**Extended Data Fig.5.**
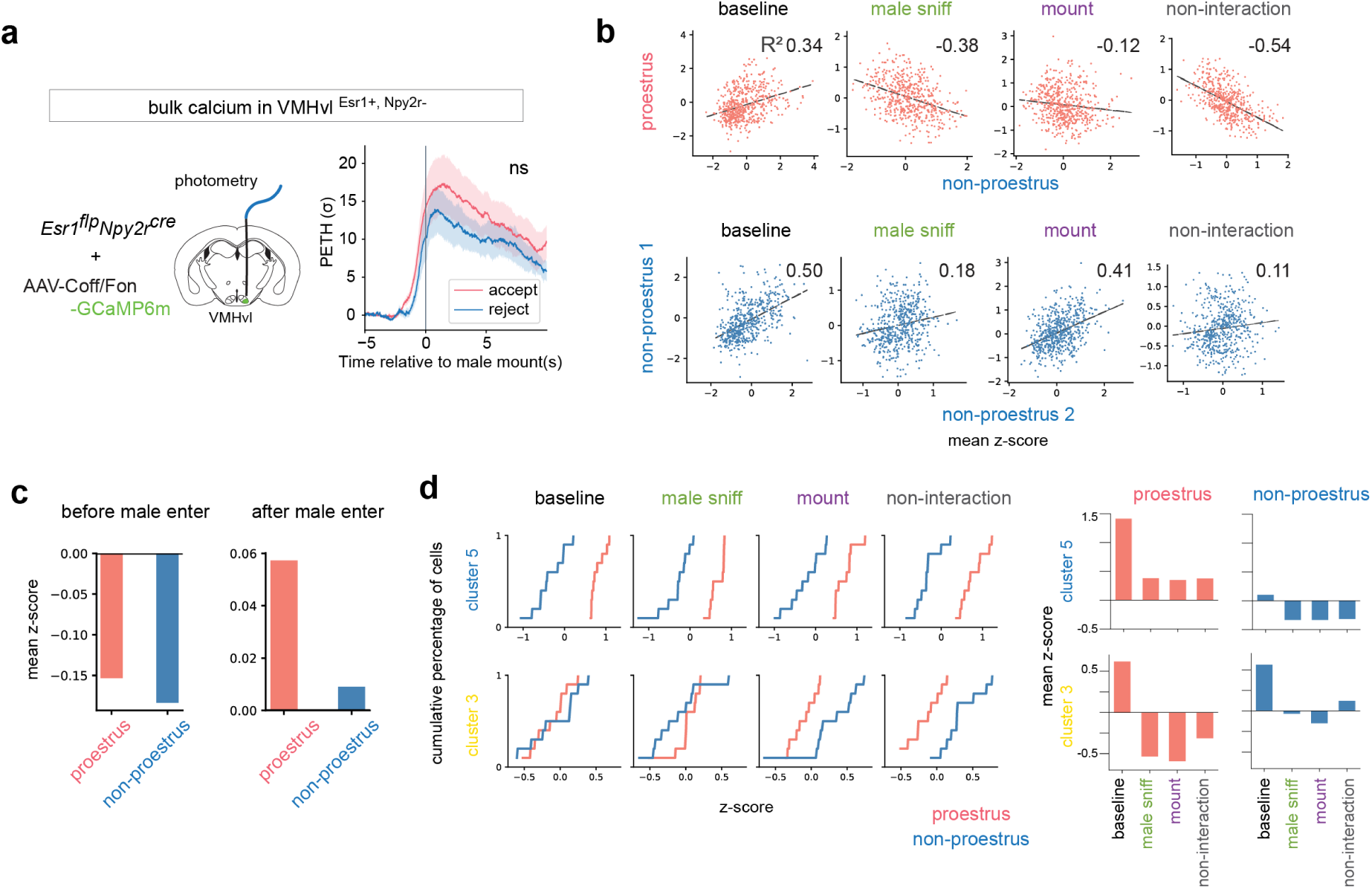
additional information for Fig.3. **a**, Photometry recording in female VMHvl α cells during accepting and rejecting mating interactions. **b,** Correlation of single unit activity during indicated behaviors between states (upper/red: proestrus vs. pooled non-proestrus; lower/blue; non-proestrus 1 vs. non-proestrus 2). Numbers indicate R^2^ value. Male sniffing and attempted mounting but not intromission (which does not occur during non-proestrus states) were compared. “non-interaction” indicates the time when the female and male are not physically interacting during freely moving mating interaction. **c,** Average activity from all cells at proestrus and non-proestrus states before and after male entry. Z-scores are calculated using ΔF/F averaged across all frames of recording period as denominator, hence negative values during pre-male baseline (left). **d,** Left, Cumulative distribution of mean z-scored neuronal activity in indicated clusters during different social behaviors. Colors indicate proestrus (red) vs. pooled non-proestrus (blue). Right, mean z-scored activity of selected clusters during indicated social behaviors.

**Extended Data Fig.6.**
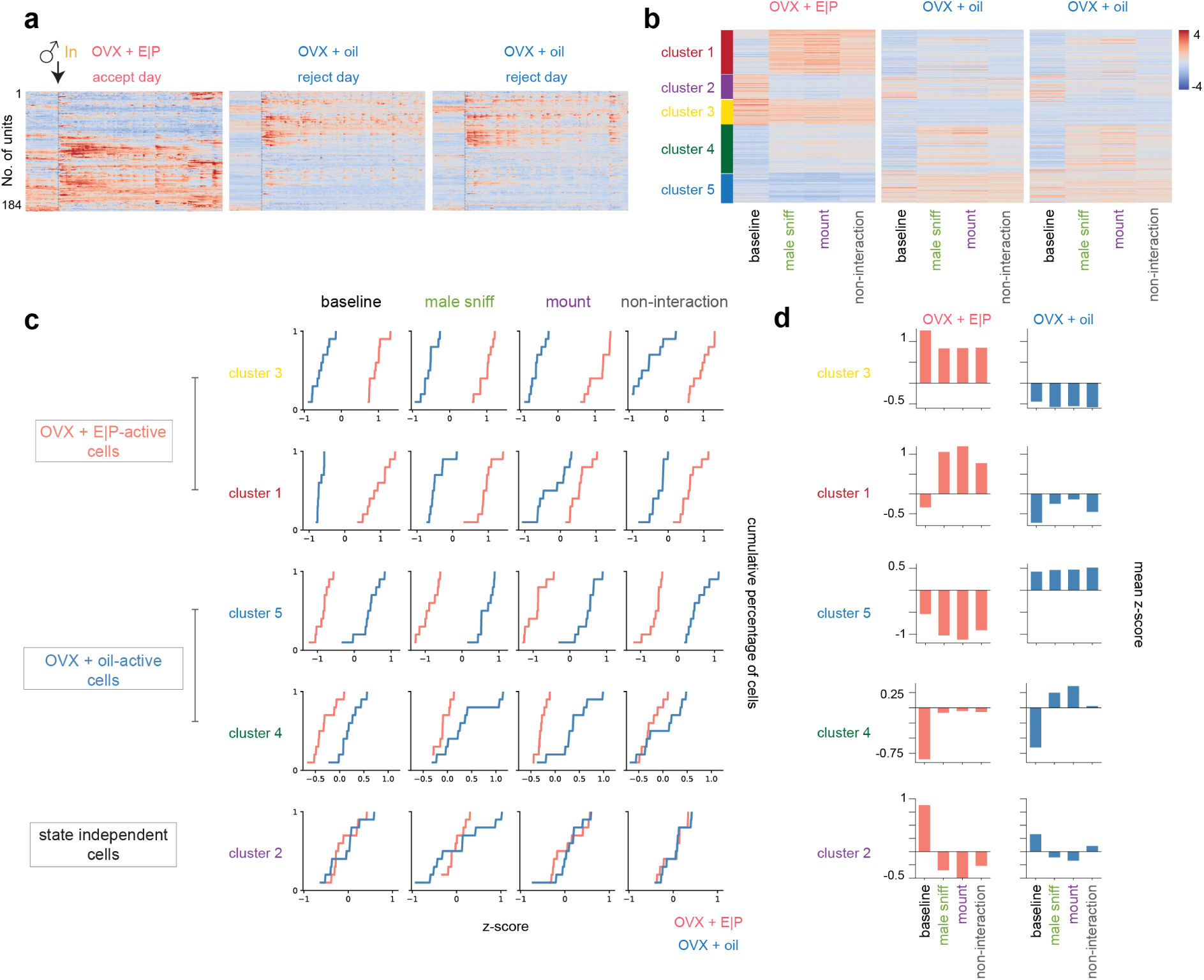
Single cell analysis for OVX females, additional information for Fig.3. **a**, Example neural traces from one ovariectomized longitudinally imaged female. **b,** Unsupervised clustering of neuronal activity in ovariectomized females during interactions with a male in hormone primed (E|P) and unprimed (“oil”) states. Combined data from N = 4 mice. Units z-scored across days. Heat map indicates standard deviations from the mean (computed across all frames). **c,** Cumulative distribution of mean neuronal activity in indicated clusters during different social behaviors. Colors indicate hormone primed (red) vs. unprimed (blue) states. **d,** Mean z-scored activity of selected clusters during indicated social behaviors.

**Extended Data Fig.7.**
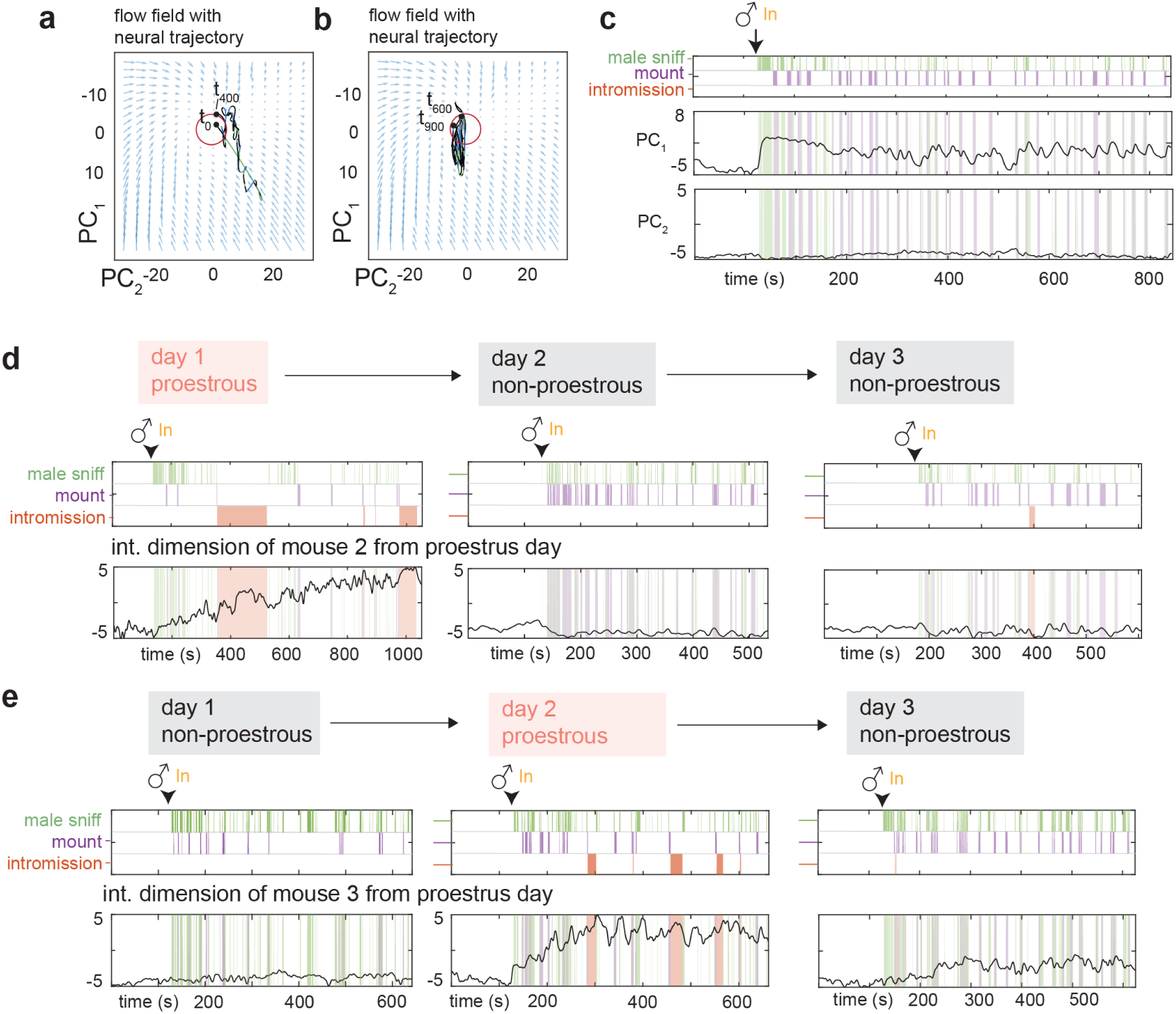
Dynamical system modelling in non-proestrus days, additional information for Fig.4. **a**, Neural trajectories and flow field of VMHvl dynamical system during non-proestrus day for t = 0 to t = 400s. **b,** Same as for t = 400s to t = 900s. **c,** Low dimensional principal components of VMHvl α dynamical system in proestrus day of mouse 1 with neural data projected from non-proestrus day. **d,** Dynamics of integration dimension in VMHvl mouse 2 discovered during proestrus day compared to activity of the same dimension on non-proestrus days. **e,** Same as **d** for mouse 3.

**Extended Data Fig.8.**
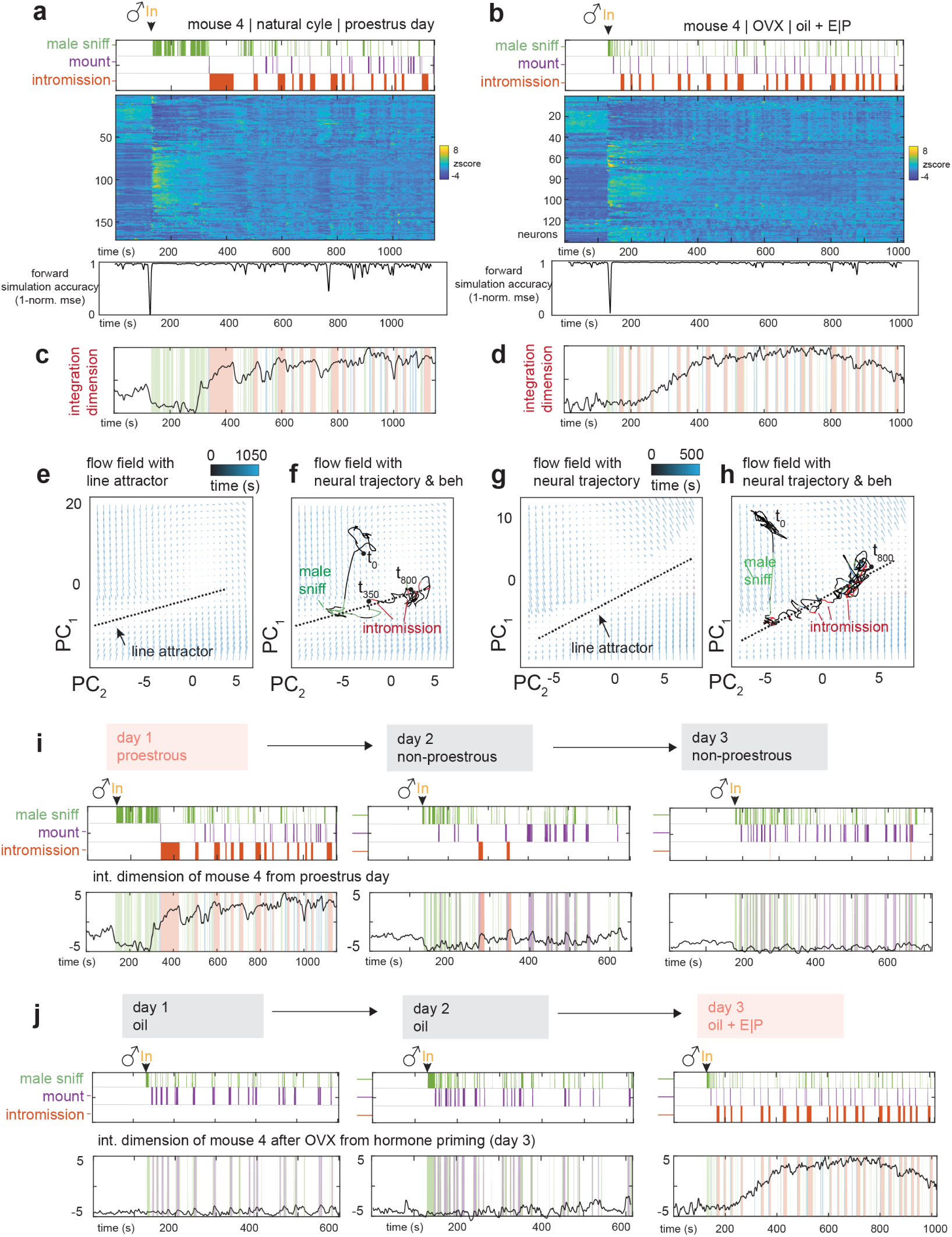
Natural cycling vs. OVX hormone priming, additional information for Fig.4. (**a, b,**) Neural raster and behaviors and rSLDS model performance (measured as forward simulation error, see Methods) for mouse 4 in proestrus day of natural estrus cycle **a,** and same mouse 4 on hormone primed day after OVX (day3, oil + E|P) **b. (c, d,)** Integration dimension identified by rSLDS on proestrus day in mouse 4 **c,** and during hormone primed day after OVX **d**. (**e, f,**) Flow field **e**, and neural trajectories of dynamical system **f**, with line attractor highlighted of model fit during the proestrus state of the estrus cycle. **(g, h,)** Same as **e, f,** for model fit during hormone primed day after OVX. **i,** Dynamics of integration dimension in mouse 4 discovered during proestrus day compared to activity of the same dimension on non-proestrus days. **j,** Dynamics of integration dimension in mouse 4 discovered during hormone primed day (day 3) compared to the activity of the same dimension during non-primed days.

**Extended Data Fig. 9.**
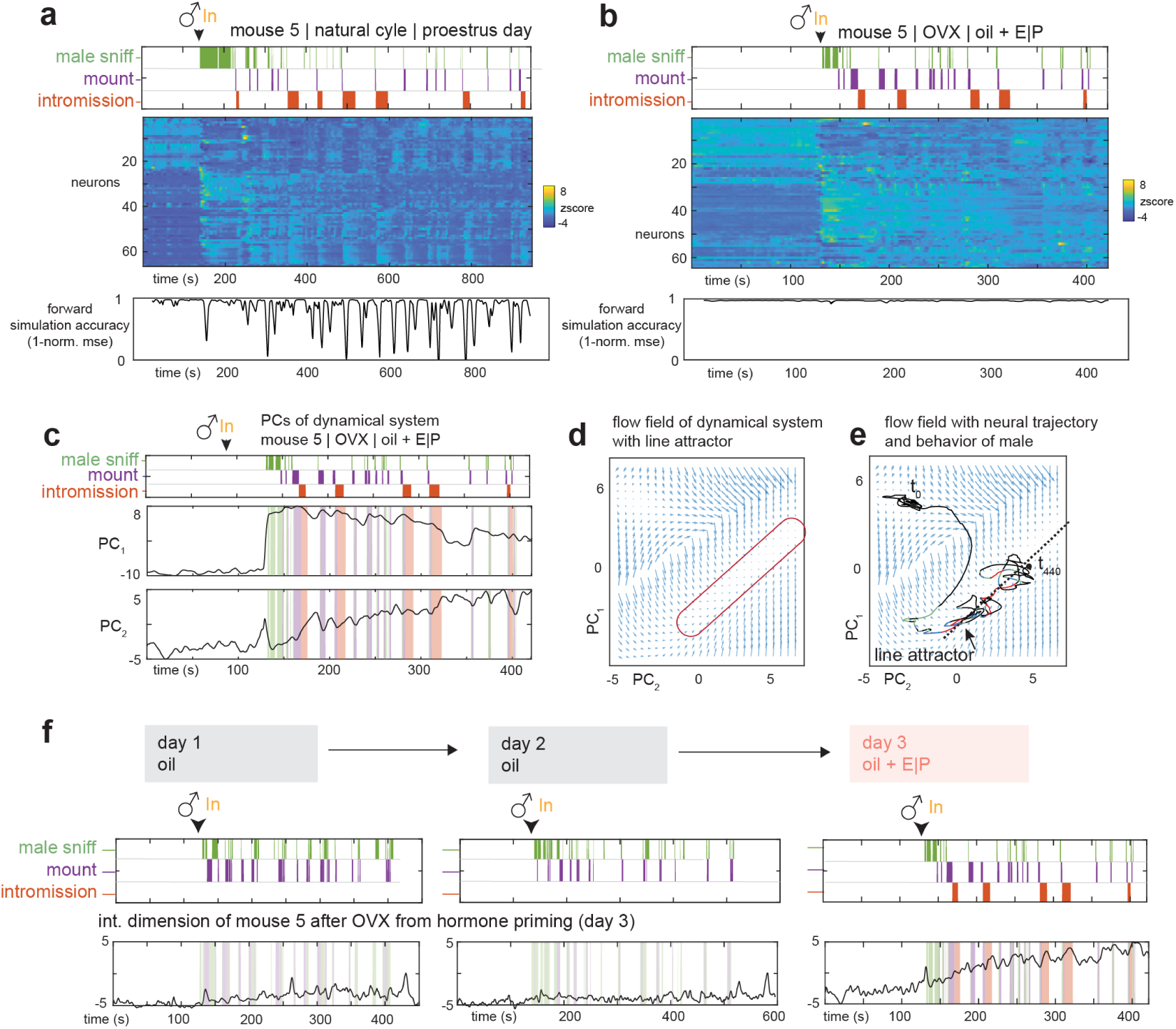
Increase of rSLDS model fits after OVX hormone priming. (**a, b,**) Neural raster and behaviors and rSLDS model performance for mouse 5 in proestrus day of natural estrus cycle **a,** and same mouse 5 on hormone primed day after OVX (day3, oil + E|P) **b. c,** Principal components of mouse 5 dynamic system fit during hormone primed day. (**d, e,**) Flow field **d**, and neural trajectories of dynamical system **e**, with line attractor highlighted of model fit during the hormone primed day after OVX in mouse 5. **f,** Dynamics of integration dimension in mouse 5 discovered during hormone primed day (day 3) compared to the activity of the same dimension during non-primed days.

**Extended Data Fig. 10.**
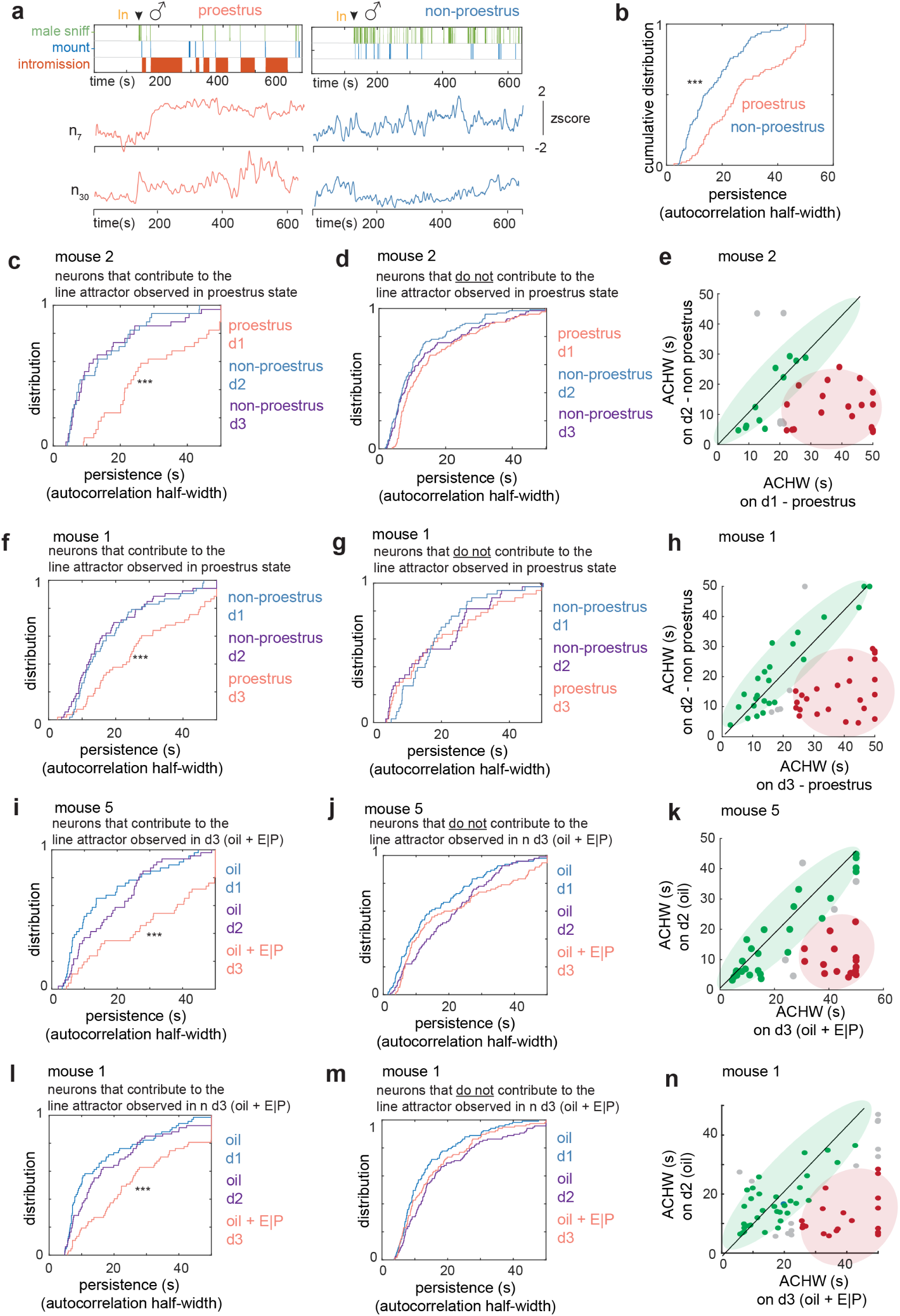
Line attractor cells display hormonal state-dependent persistence. **a**, Example units active during both proestrus (red traces, left) and non-proestrus (blue traces, right), showing persistence on proestrus day and fast dynamics on the non-proestrus days. **b,** Comparison of cumulative distribution of ACHWs to that of same neurons on non-proestrus days. Data from mouse 1. **c,** Cumulative distribution of ACHWs for units with significant weights on integration dimension across proestrus and non-proestrus day. Data from mouse 1. **d,** Cumulative distribution of ACHWs for mouse 1, for units that do not contribute to the integration dimension on the proestrus day, compared on proestrus vs non-proestrus days. **e,** Scatter plot of ACHWs for units with significant weights on integration dimension for proestrus day vs non-proestrus day. Data from mouse 1. (**f-h**) Same as **c-e** for mouse 2. **i,** Cumulative distribution of ACHWs for units with significant weights on integration dimension across hormone primed (day 3) and non-primed days (days 2, 1). Data from mouse 5. **j,** Cumulative distribution of AHWs for mouse 1, for units that do not contribute to the integration dimension across hormone primed (day 3) and non-primed days (days 2, 1). Data from mouse 5. **k,** Scatter plot of ACHWs for units with significant weights on integration dimension for hormone-primed day vs non-primed day. Data from mouse 1. (**l-n**) Same as **i-j** for mouse 1.

## Methods

### Mice

All experimental procedures involving the use of live mice or their tissues were carried out in accordance with NIH guidelines and approved by the Institute Animal Care and Use Committee (IACUC) and the Institute Biosafety Committee (IBC) at the California Institute of Technology (Caltech). All mice in this study, including wild-type and transgenic mice, were bred at Caltech or purchased from Charles River Laboratory. Singly housed C57BL/6N female mice (2-5 months) were used as experimental mice. *Npy2r^cre^* mice (Jackson Laboratory stock no. 029285) (=N1), *Esr1^cre^* mice, *Esr1^flp^* mice (>N10) were backcrossed into the C57BL/6N background and bred at Caltech. Heterozygous *Npy2r^cre^*, *Esr1^cre^* or double heterozygote *Esr1^flp/+^Npy2r^cre/+^* mice were used for cell-specific targeting experiments and were genotyped by PCR analysis using genomic DNA from tail tissue. All mice were housed in ventilated micro-isolator cages in a temperature-controlled environment (median temperature 23°C, humidity 60%), under a reversed 11-h dark–13-h light cycle, with ad libitum access to food and water. Mouse cages were changed weekly.

### Surgeries

Surgeries were performed on female Esr1^Flp/+^Npy2r ^Cre/+^ females aged 2 months. Virus injection and implantation were performed as described previously^14, 20^. Briefly, animals were anaesthetized with isoflurane (5% for induction and 1.5% for maintenance) and placed on a stereotaxic frame (David Kopf Instruments). Virus was injected into the target area using a pulled-glass capillary (World Precision Instruments) and a pressure injector (Micro4 controller, World Precision Instruments), at a flow rate of 20 nl min^-1^. The glass capillary was left in place for 10 min following injection before withdrawal. Lenses were slowly lowered into the brain and fixed to the skull with dental cement (Metabond, Parkell). Females were co-housed with a vasectomized male mouse after virus injection and lens implantation. Four weeks after lens implantation, mice were head-fixed on a running wheel and a miniaturized micro-endoscope (nVista, Inscopix) was lowered over the implanted lens until GCaMP-expressing fluorescent neurons were in focus. Once GCaMP-expressing neurons were detected, a permanent baseplate was attached to the skull with dental cement. The co-housed vasectomized males were removed.

### Virus injection and GRIN lens implantation

The following AAVs were used in this study, with injection titers as indicated. Viruses with a high original titer were diluted with clean PBS on the day of use. AAV-DJ-EF1a-Coff/Fon-GCaMP6m (4.5 x e12, Addgene plasmid) was packaged at the HHMI Janelia Research Campus virus facility. “Coff/Fon” indicates Cre-OFF/FLP-ON virus. Stereotaxic injection coordinates were based on the Paxinos and Franklin atlas. Virus injection: VMHvl, AP: −1.6, ML: ±0.78, DV: −5.73; GRIN lens implantation: VMHvl: AP: −1.6, ML: −0.75, DV: −5.6 (⌀0.6 × 7.3 mm GRIN lens).

### Vaginal cytology

To determine the estrus phases of tested females, vaginal smear cytology was applied on the same day as the behavior test. A vaginal smear was collected immediately after the behavioral test and stained with 0.1% crystal violet solution for 1 minute. Cell types in the stained vaginal smear were checked microscopically. In this study, the proestrus phase was characterized by many nucleated epithelial, some cornified epithelial and no leukocytes.

### Hormone priming

Female mice were ovariectomized and estrus was induced by hormone priming. Estradiol benzoate (E2) and progesterone (PG) powder was dissolved in sesame oil. For primed females, 50ul 200ng/ml E2 was delivered subcutaneously on day-2 and day-1 at 3pm. 10mg/ml PG was delivered subcutaneously on the day of test at 10am. Behavior test was performed 4-6 hours after PG injection. For unprimed female, sesame oil was injected at the same time points as hormone injections. Vaginal smear cytology was applied on the same day as the behavior test to make sure that the females were completely primed or unprimed.

### Sex representation assay

All behavior tests were performed under red light. Group housed C57BL/6N male and female mice (2-4 months) were used for the test. Tested female was acclimated in her home cage under the recording setup^28^ for 10 minutes. A toy, a female or a male was introduced to the tested female with a 90-second interval. Each interaction lasted for 1 minute. The sequential representations were repeated for 3 times.

### Mating assay

Singly housed sexually experienced C57BL/6N male were used for mating assay. Male mice used for test were initially co-housed with a female mouse for at least 1 week and singly housed at least 1 week before test. On the day of test. Male mouse was acclimated in his home cage under the recording setup. A random female mouse was placed into male cage until three male mounting bouts were observed. The tested female mice were acclimated in a new cage for 10 minutes before being introduced into the male cage. The male contact mating interaction lasted for 5-15 mins. At the end of the free interaction, a pencil cup was introduced to restrain the male. Then the imaging and behavior recording during the non-contact period continued for 3-5 mins.

### Behavior annotations

Behavior videos were manually annotated using a custom MATLAB-based behavior annotation interface. A’baseline’ period of 2 min when the animal was alone in its cage was recorded at the start of every recording session. In the male contact interaction assay, five behaviors were annotated: baseline (10-20 seconds before female was introduced into male cage), male sniff (male sniff female with physical contact), mount, intromission, no-interaction (periods when the animals were not physically interacting). During the non-contact interaction, female sniff (female sniff the pencil cup with male inside) was annotated.

### Fiber photometry

The fiber photometry setup was as previously described in earlier research with minor modifications. We used 470 nm LEDs (M470F3, Thorlabs, filtered with 470-10 nm bandpass filters FB470-10, Thorlabs) for fluorophore excitation, and 405 nm LEDs for isosbestic excitation (M405FP1, Thorlabs, filtered with 410–10 nm bandpass filters FB410-10, Thorlabs). LEDs were modulated at 208 Hz (470 nm) and 333 Hz (405 nm) and controlled by a real-time processor (RZ5P, Tucker David Technologies) via an LED driver (DC4104, Thorlabs). The emission signal from the 470 nm excitation was normalized to the emission signal from the isosbestic excitation (405 nm), to control for motion artefacts, photobleaching and levels of GCaMP6m expression. LEDs were coupled to a 425 nm longpass dichroic mirror (Thorlabs, DMLP425R) via fiber optic patch cables (diameter 400 mm, N.A., 0.48; Doric lenses). Emitted light was collected via the patch cable, coupled to a 490 nm longpass dichroic mirror (DMLP490R, Thorlabs), filtered (FF01-542/27-25, Sem-rock), collimated through a focusing lens (F671SMA-405, Thorlabs) and detected by the photodetectors (Model 2151, Newport). Recordings were acquired using Synapse software (Tucker Davis Technologies). On the test day, after at least 5 minutes of acclimation under the recording setup, the female was first recorded for 5 minutes to establish a baseline. Then behavior assays were proceeded and fluorescence were recorded for the indicated period of time, as described in the text. All data analyses were performed in Python. Behavioral video files and fiber photometry data were time-locked. Fn was calculated using normalized (405 nm) fluorescence signals from 470 nm excitation. Fn(t) = 100 x [F470(t) – F405fit(t)] / F405fit(t). For the peri-event time histogram (PETH), the baseline value F_0_ and standard deviation SD_0_. was calculated using a-5 to-3 second window. Overlapping behavioral bouts within this time window were excluded from the analysis. Then PETH was calculated by [(Fn(t) – F_0_]/SD_0_.

### Micro-endoscopic imaging data Acquisition

Imaging data were acquired at 30 Hz with 2× spatial downsampling; light-emitting diode power (0.2–0.5) and gain (1–8×) were adjusted depending on the brightness of GCaMP expression as determined by the image histogram according to the user manual. A transistor–transistor logic (TTL) pulse from the Sync port of the data acquisition box (DAQ, Inscopix) was used for synchronous triggering of StreamPix7 (Norpix) for video recording.

### Micro-endoscopic data extraction and preprocessing

Miniscope data were acquired at 30 Hz using the Inscopix Data Acquisition Software as 2× down sampled.isxd files. Preprocessing and motion correction were performed using Inscopix Data Processing Software. Briefly, raw imaging data from three recording dates were concatenated. 2× spatially down sampled, motion corrected and temporally down sampled to 10 Hz. Further and exported as a.tiff image stack. A spatial band-pass filter was then applied to remove out-of-focus background. After preprocessing, calcium traces were extracted and deconvolved using the CNMF-E large data pipeline with the following parameters: patch_dims = [32, 32], gSig = 3, gSiz = 13, ring_radius = 19, min_corr = 0.72, min_pnr = 8. The spatial and temporal components of every extracted unit were carefully inspected manually (SNR, PNR, size, motion artefacts, decay kinetics and so on) and outliers (obvious deviations from the normal distribution) were discarded. The extracted traces were then z-scored before analysis.

### Longitudinal imaging data extraction and preprocessing

The females performed mating assay and were imaged for consecutive 3-5 days. Miniscope data from one proestrus accepting day and two non-proestrus rejecting days were selected and concatenated to one.isxd file. Data was preprocessed and the traces were extracted as described in the last section. The three-day concatenated traces were z-scored, and then split to multiple traces for individual days.

### Choice probability

Choice probability is a metric that estimates how well either of two different behaviors can be predicted/distinguished, based on the activity of any given neuron during these two behaviors. CP of single neurons was computed using previously described methods^20^. To compute the CP of a single neuron for any behavior pair, 1 s binned neuronal responses occurring during each of the two behaviors were used to generate a receiver operating characteristic curve. CP is defined as the area under the curve bounded between 0 and 1. A CP of 0.5 indicates that the activity of the neuron cannot distinguish between the two alternative behaviors. We defined a neuron as being capable of distinguishing between two behaviors if the CP of that neuron was >0.7 or <0.3 and was >2 σ or <−2 σ of the CP computed using shuffled data (repeated 1000 times).

### Unsupervised clustering

A *k*-means clustering algorithm was used for unsupervised clustering. The number of *k*=5 was determined using the Elbow method and Silhouette method. For Fig. 3, clustering was performed on data from the proestrus day, then the cluster ids were assigned to the same units in the other two non-proestrus days.

### Generalized linear model

To predict neural activity from behavior, we trained generalized linear models to predict the activity of each neurons k, as a weighted linear combination of 3 male behaviors: male sniffing, mounting and intromission as follows:

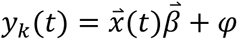

Here, *y_k_*(*t*) is the calcium activity of neuron k at time t, 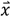(t) is a matrix of 3 male behaviors provided as binarized vectors. 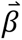 is a behavior-filter which described how a neuron integrates stimulus over a 10s period (example filters are shown in Extended Data Fig.1d-e). *φ* is an error term. The model was fit using 10-fold cross validation and model performance is reported as cross-validated *R*^2^ (cv*R*^2^).

### Dynamical system modelling

Recurrent-switching linear dynamical system (rSLDS) models^16, 29^ are fit to neural data as previously described^15^. Briefly, rSLDS is a generative state-space model that decomposes non-linear time series data into a set of linear dynamical systems, also called ‘states’. The model describes three sets of variables: a set of discrete states (z), a set of latent factors (x) that captures the low-dimensional nature of neural activity, and the activity of recorded neurons (y). While the model can also allow for the incorporation of external inputs based on behavior features, such external inputs were not included in our first analysis.

The model is formulated as follows: At each time t = 1,2,…T_n_, there is a discrete state *Z_t_* ∈ {1,2,…,*K*}. In rSLDS. These states depend recurrently on the continuous latent factors (x) as follows:

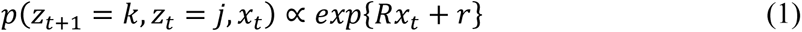

where *R* and *r* parameterizes a map from the previous discrete state and continuous state using a softmax link function to a distribution over the next discrete states. The discrete state *Z*_t_ determines the linear dynamical system used to generate the latent factors at any time t:

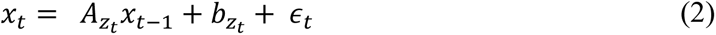

where *A_k_* ∈ ℝ^*d*×*d*^ is a dynamics matrix and *b_k_* ∈ ℝ^*d*^ is a bias vector, where *d* is the dimensionality of the latent space and *ϵ_t_* ∼ *N*(0,*Q_zt_*) is a Gaussian-distributed noise (aka innovation) term.

Lastly, we can recover the activity of recorded neurons by modelling activity as a linear noisy Gaussian observation *y_t_* ∈ ℝ*^N^* where *N* is the number of recorded neurons:

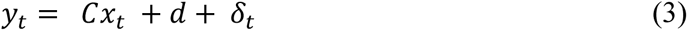

For *C* ∈ ℝ^*N*×*D*^ and *δ_t_* ∼ *N*(0,*S*), a Gaussian noise term. Overall, the system parameters that rSLDS needs to learn consists of the state transition dynamics, library of linear dynamical system matrices and neuron-specific emission parameters, which we write as:

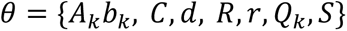

We evaluate model performance using both the evidence lower bound (ELBO) and the forward simulation accuracy (FSA) (Fig. 3a) described in Nair et al., 2023^15^ as well as by calculating the variance explained by the model on data. Briefly, given observed neural activity in the reduced neural state space at time t, we predict the trajectory of population activity over an ensuing small time interval Δt using the fit rSLDS model, then compute the mean squared error (MSE) between that trajectory and the observed data at time t+ Δt. This MSE is computed across all dimensions of the reduced latent space and repeated for all times t across cross validation folds. This error metric is normalized to a 0-1 range in each animal across the whole recording to obtain a bounded measure of model performance. The FSA can intuition of where model performance drops during the recording. In addition to MSE, we also calculate the Pearson’s correlation coefficient (*R^2^*) between the predicted and observed data for each dimension following the forward simulation. By taking the average correlation coefficient across dimensions, we can obtain a quantitative estimate of variance explained by rSLDS on observed data.

The number of states and dimensions used for the model are determined using 5-fold cross validation. Visualization of the dynamical system using principal components analysis is performed as described previously ^15^.

In addition to fitting models without inputs, we also refit the same models for all animals using behaviors performed by the male intruder as inputs including male sniffing, mounting and intromission. In this formulation, equations (1) and (2) which describe discrete state and latent factor dynamics are modified to include an input term that describes how male behaviors can contribute to state transitions and neural dynamics. This is formulated as:

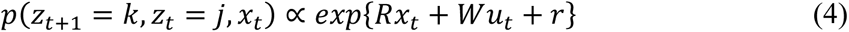

where R, W and r parameterizes a map from the previous discrete state (z), latent factors (x) and external inputs (u) using a softmax link function to a distribution over the next discrete states. In a given discrete state, latent factors (x) now evolve as:

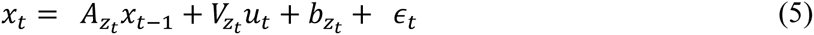

where *A_k_* is the dynamics matrix, *v_k_* describes the influence of external inputs to ongoing dynamics and *b_k_* is a bias vector. These external inputs also influence the neural data directly as follows:

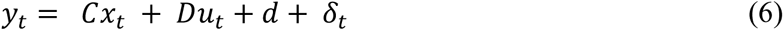

where *C* ∈ ℝ^N×D^, *D* ∈ ℝ^N×M^ where M is the dimensionality of the external inputs and *δ_t_* ∼ *N*(0,*S*), a Gaussian noise term. In this formulation, the rSLDS strictly generalizes the GLM described above.

These new parameters are estimated using variational expectation-maximization as mentioned previously and the fit quality is assessed using the forward simulation accuracy metric and variance explained.

Code used to fit rSLDS on neural data is available in the SSM package: (https://github.com/lindermanlab/ssm)

### Estimation of time constants

We estimated the time constant of each dimension of linear dynamical systems using eigenvalues *λ_a_* of the dynamics matrix of that system, derived previously^30^ as:

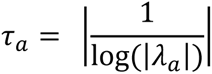

### Calculation of auto-correlation half-width

We computed autocorrelation halfwidths by calculating the autocorrelation function for each neuron timeseries data (y_t_) for a set of lags as described previously^12^. Briefly, for a time series (y_t_), the autocorrelation for lag k is:

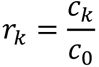

where *c_k_* is defined as:

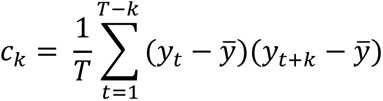

and *c*_0_ is the sample variance of the data. The half-width is found for each neuron as the point where the autocorrelation function reaches a value of 0.5 (Extended Data Fig.4d-e).

## Data availability

The data on which this study is based are available on reasonable request.

## Code availability

The custom MATLAB and Python codes used to analyse the data in this study are available on request.

## Acknowledgments

We thank A. Kennedy and S. Linderman for critical feedback on the manuscript, Y. Huang for genotyping, G. Mancuso for administrative assistance, C. Chiu for lab management, E. Carcamo for mouse colony management, Caltech OLAR staff for animal care, and members of the Anderson Laboratory for helpful comments on this project. DJA is an investigator of the Howard Hughes Medical Institute. This work was supported by grants from the NIH (RO1MH112593, RO1MH123612 and RO1NS123916). A.N is supported by a National Science Scholarship from the Agency of Science, Technology and Research, Singapore.

## Author contributions

Conceptualization: D.J.A., M.L.; Experiments: M.L.; Data analysis: M.L., A.N., S.L.; Supervision: D.J.A.; Writing: D.J.A., M.L., A.N.

