## Supplementary_information for "Periodic hypothalamic attractor-like dynamics during the estrus cycle"

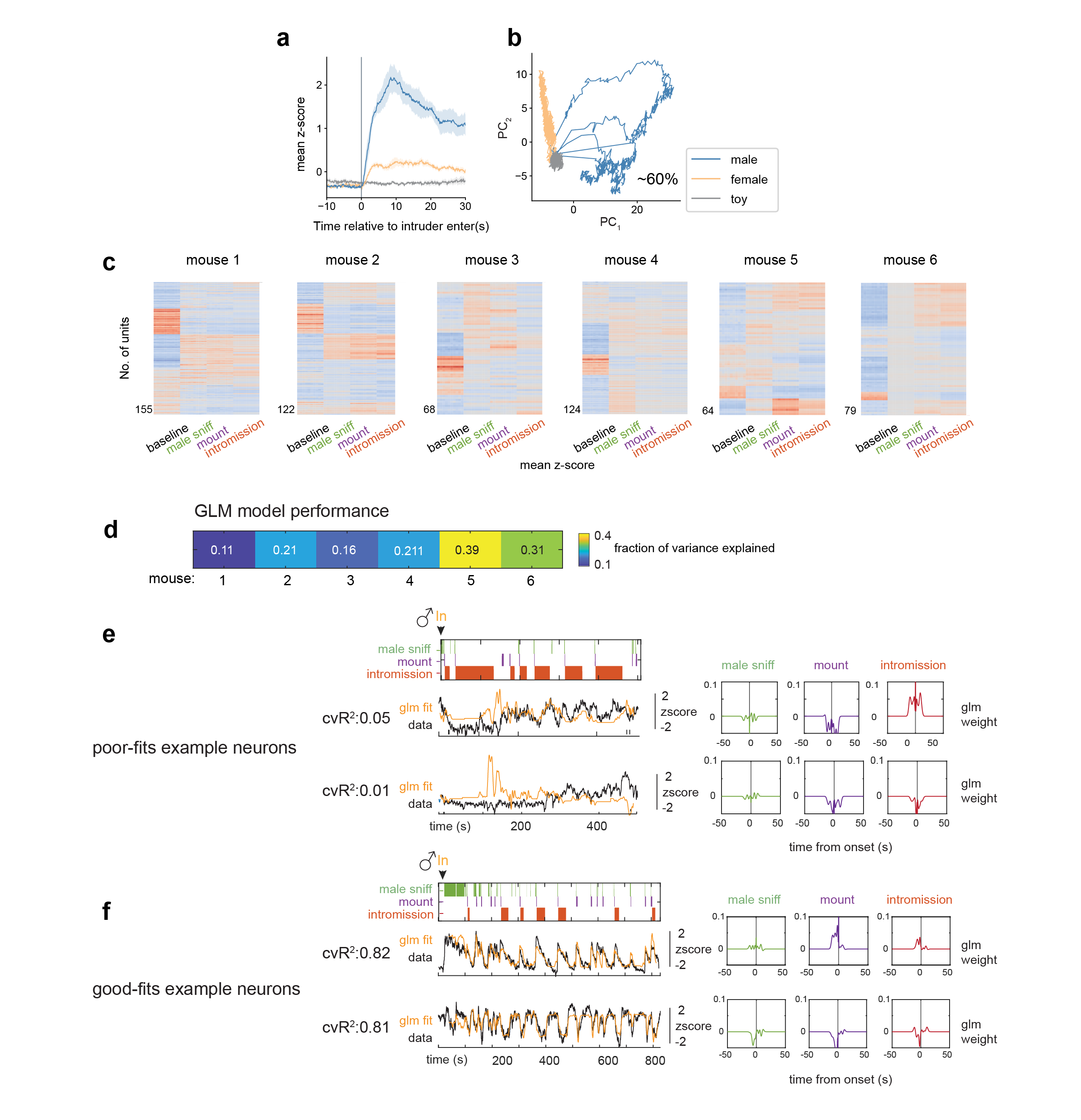
**Extended Data Fig.1. | Additional information for Fig. 1. a,** Mean responses of female VMHvl^Esr1^ α cells to male, female and toy. N=8 mice. **b,** PCA of neuronal responses to male, female and toy from one example female. **c,** Unsupervised clustering of neuronal activity from individual mice. Columns represents activity averaged across 1s windows during each indicated behavior. **d,** GLM model performance (mean fraction of variance explained across all neurons) in each mouse. **e,f,** Example generalized linear model fits and behavior filters for poorly fit neurons **e**, from mouse 1 and well fit neurons **f**, from mouse 5.

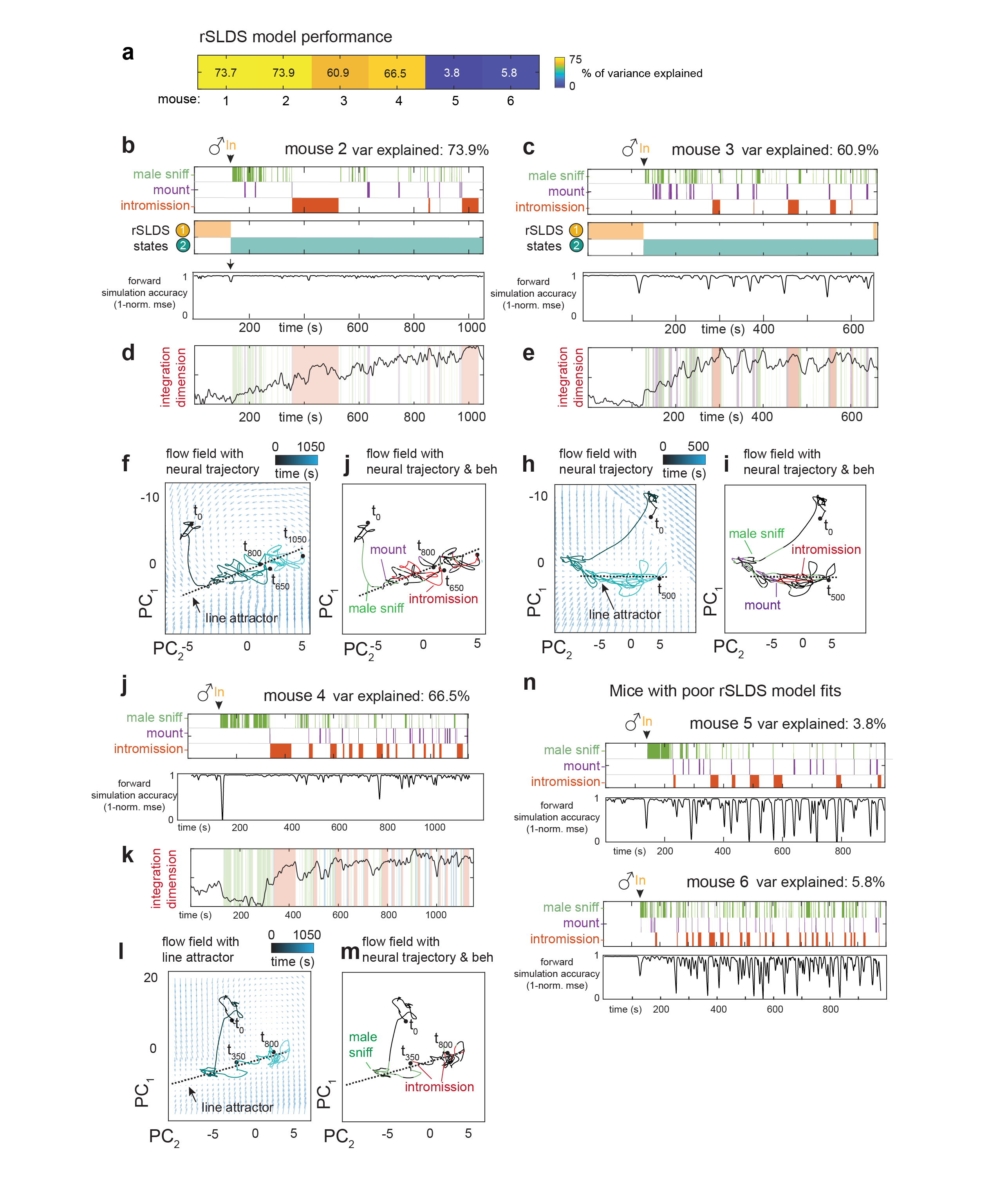
**Extended Data Fig.2 | rSLDS model fitting for all mice, additional information for Fig.2. a,** rSLDS model performance (mean fraction of variance explained across all dimensions in given mouse) in each mouse. **b,** States discovered by recurrent switching linear dynamical systems (rSLDS) aligned to behaviors performed by the male intruder towards female mouse 2 (Top). rSLDS model fit (forward simulation accuracy, bottom). **c,** Same as **b** for mouse 3. **d,** Dynamics of integration dimension in mouse 2. **e,** Same as **d** for mouse 3. **f,** Flow field of VMHvl α dynamical system for mouse 2 showing neural trajectories in state space, annotated by time from male encounter (t_0_). **g,** Neural state space of VMHvl α dynamical system for mouse 2 highlighting male behaviors and region containing approximate line attractor. **h,** Same as **f** for mouse 3. **i,** Same as **g** for mouse 3. **j,** rSLDS model fit for mouse 4. **k,** Dynamics of integration dimension in mouse 4. **l,** Same as **f** for mouse 4. **m,** Same as **g** for mouse 4. **n,** rSLDS model fit for mouse 5 and 6 highlighting a poor model fit in these animals.

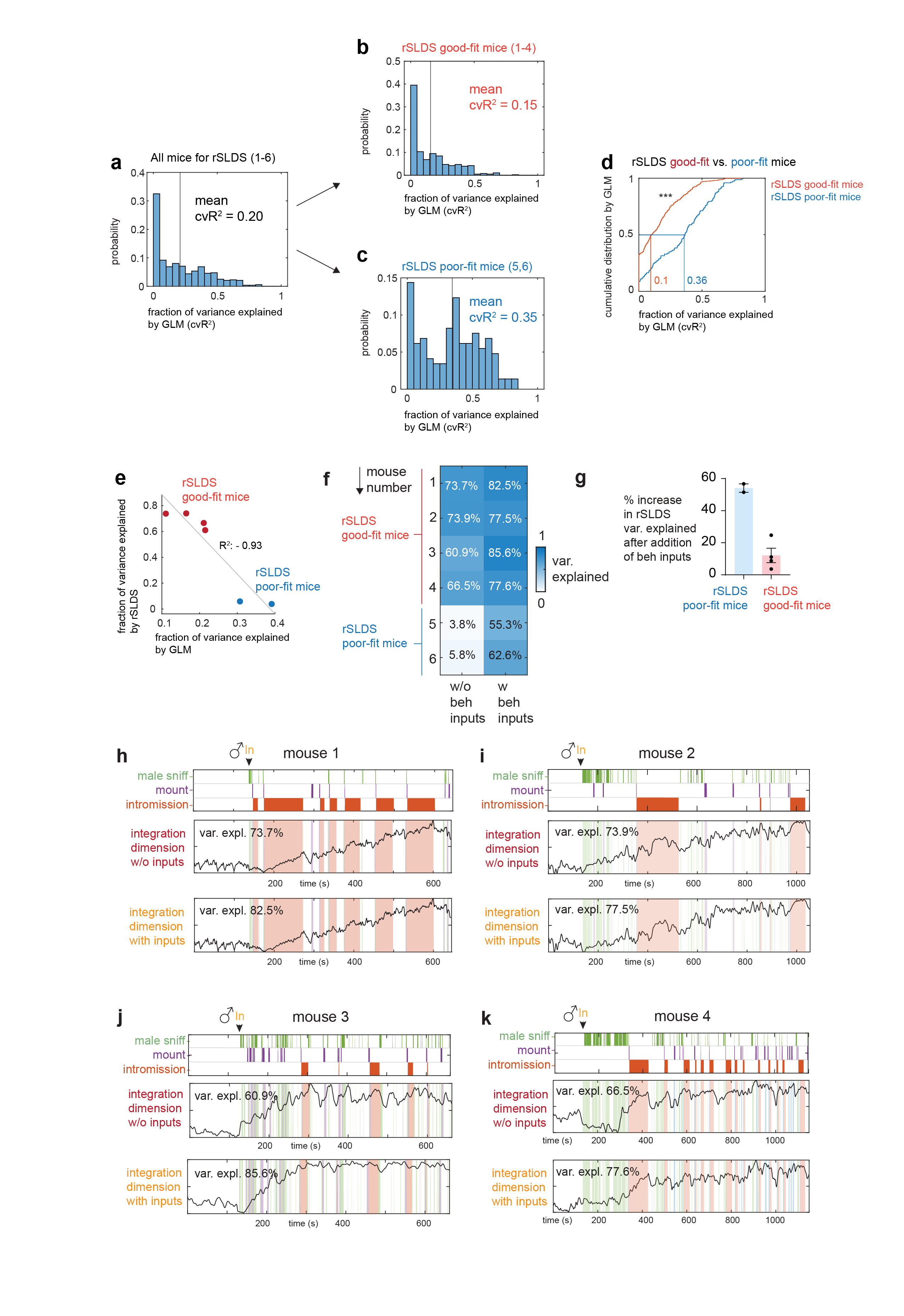
**Extended Data Fig.3 | GLM fits for rSLDS good-fit and poor-fit mice, additional information for Fig.2. a,** Variance explained by GLM trained to predict neural activity from behavior for all mice used for rSLDS (distribution mean = 0.20, N = 6 mice). **b,** Variance explained by GLM trained to predict neural activity from behavior for mice with good rSLDS fits (distribution mean = 0.15, N = 4 mice, mouse 1-4). **c,** Variance explained by GLM trained to predict neural activity from behavior for mice with poor rSLDS fits (distribution mean = 0.35, N = 2 mice, mouse 5, 6). **d,** Cumulative distribution of GLM model fit (cross validation R^2^) for rSLDS good-fit mice (median: 0.1) vs rSLDS poor-fit mice (median: 0.36), ***p<0.0001. **e,** Correlation between variance explained by GLM and rSLDS. **f,** Variance explained by rSLDS fitting with or without behavior inputs. **g,** Percentage of increase in rSLDS variance explained after adding behavior inputs. **h,** rSLDS integration dimension in models fit with and without male behavior inputs in mouse 1. The rSLDS model performance as variance explained is also shown for each model. Percentage of increase in rSLDS variance explained after adding behavior inputs. **i,** Same as **h,** for mouse 2. **j,** Same as **h** for mouse 3. **k,** Same as **h** for mouse 4.

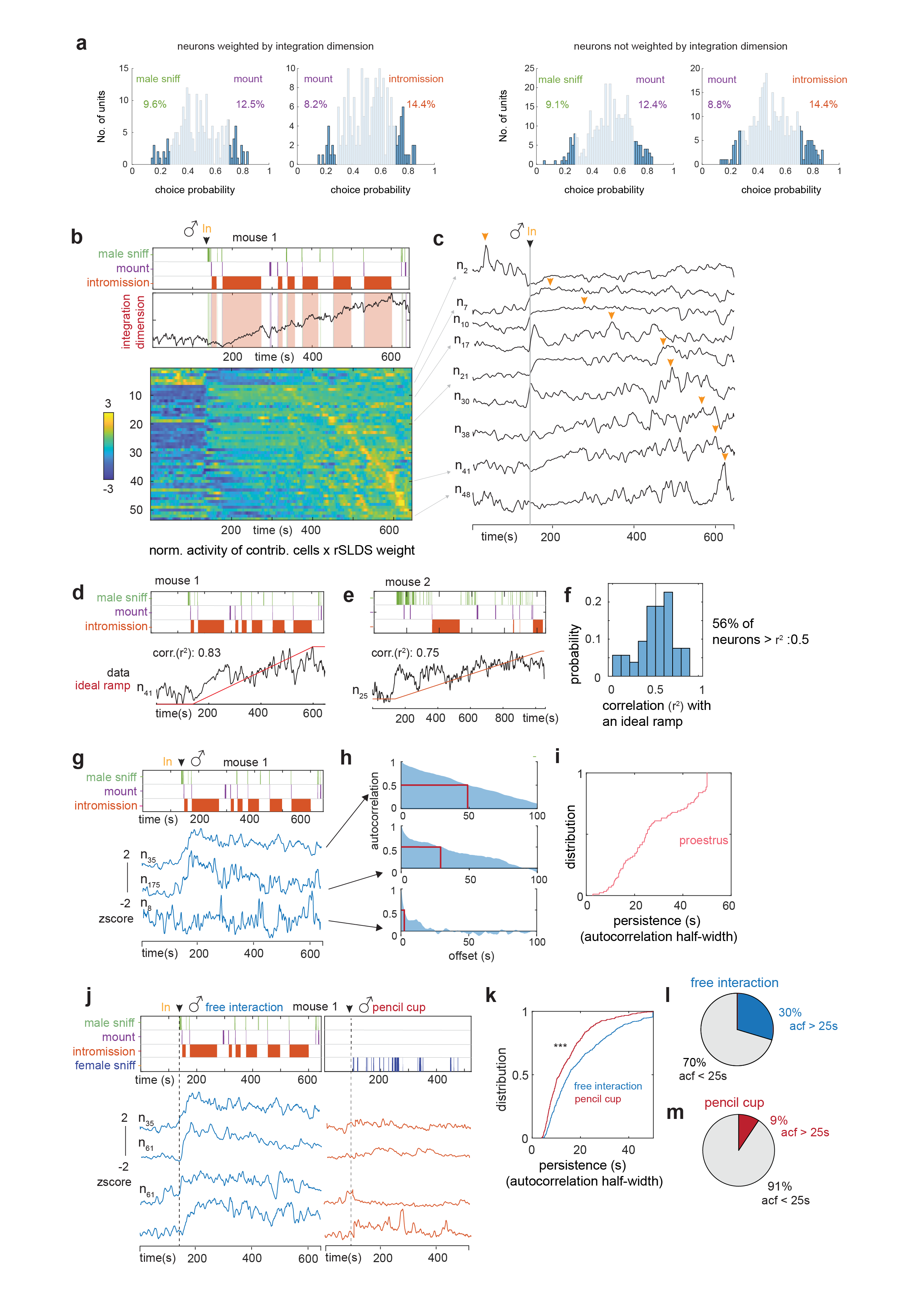
**Extended Data Fig.4 | Properties of line attractor cells, additional information for Fig.2. a,** Choice Probability (CP) histograms and number of tuned cells from highly weighted (left) cells or not highly weighted cells (right) to integration dimension. cutoff: CP>0.7 or <0.3 and >2σ. N=4 mice **b, *Upper***, relationship of male behavior to weighted average of all units contributing to integration dimension as a function of time. Data from 1 example mouse. ***Lower***, normalized activity (z-score) of individual units times rSLDS weight for each unit exhibiting a significant weight in the integration dimension of VMHvl mouse 1, sorted by time to peak. **c,** Traces of example units from **b**, *lower*. Yellow arrow indicates peak of activity for each unit. **d,** Correlation of example units activity with an ideal ramp in mouse 1. **e,** Same as D for example unit in mouse 2. **f,** Distribution of correlation of individual neuron activity with ideal ramp. **g,** Example VMHvl neurons in proestrus day showing a range of persistent activity (z-scored ∆F/F). **h,** Auto-correlation half-width (ACHW) as a measure of persistent activity, for example units shown in **g**. **i,** Cumulative distribution of ACHW for units with significant weights on the integration dimension during proestrus day (N = 4 mice). **j,** Dynamics of persistently active neurons identified during proestrus day during pencil-cup assay. **k,** Cumulative distribution of ACHW for same neurons during free interaction vs pencil cup assay ***p<0.001. **l,** Pie chart indicating fraction of neurons with ACHW > 25s in free interaction. **m,** Same as **l** for data obtained during pencil cup assay.

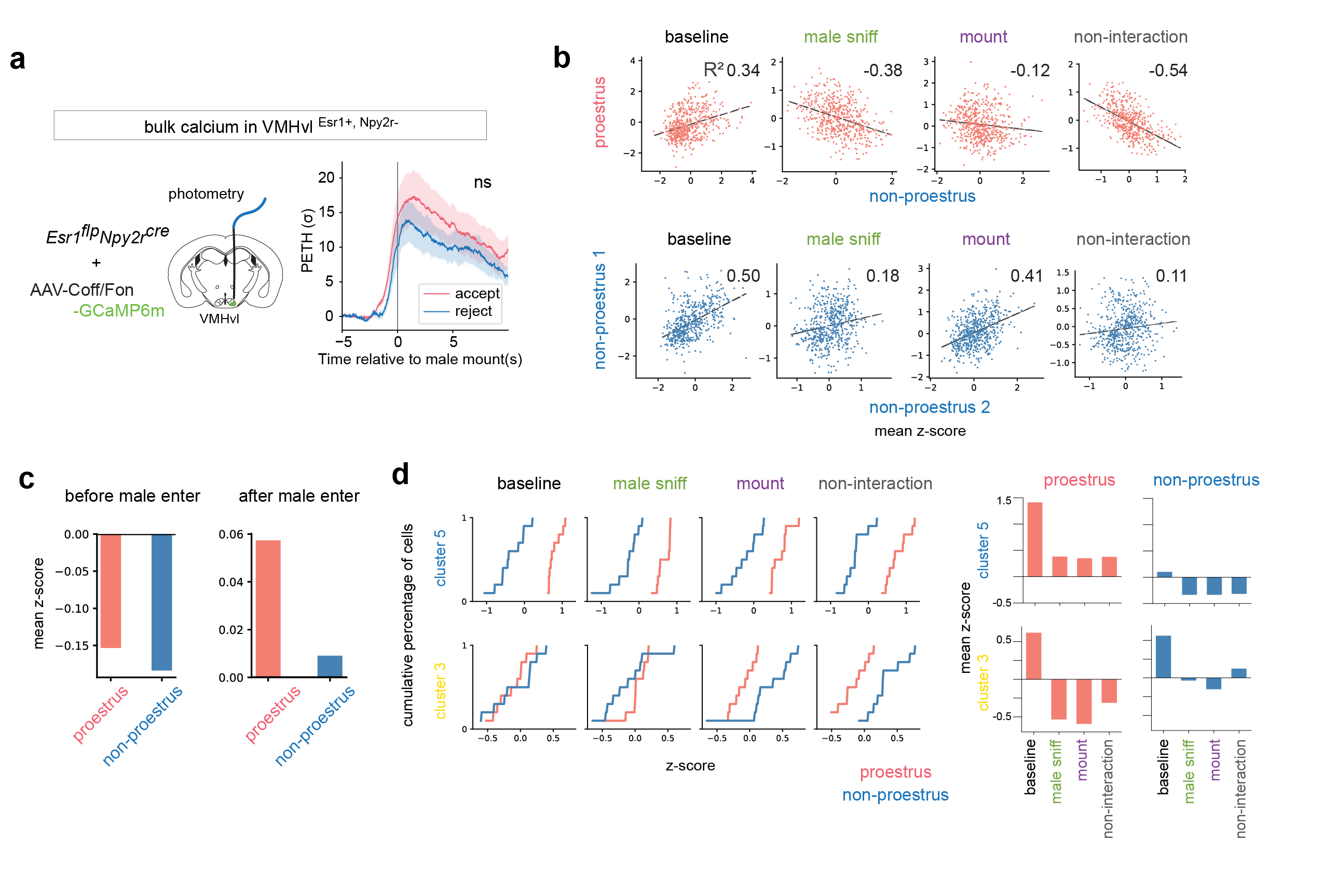
**Extended Data Fig.5 | additional information for Fig.3. a,** Photometry recording in female VMHvl α cells during accepting and rejecting mating interactions. **b,** Correlation of single unit activity during indicated behaviors between states (upper/red: proestrus vs. pooled non-proestrus; lower/blue; non-proestrus 1 vs. non-proestrus 2). Numbers indicate R^2^ value. Male sniffing and attempted mounting but not intromission (which does not occur during non-proestrus states) were compared. “non-interaction” indicates the time when the female and male are not physically interacting during freely moving mating interaction. **c,** Average activity from all cells at proestrus and non-proestrus states before and after male entry. Z-scores are calculated using ∆F/F averaged across all frames of recording period as denominator, hence negative values during pre-male baseline (left). **d,** Left, Cumulative distribution of mean z-scored neuronal activity in indicated clusters during different social behaviors. Colors indicate proestrus (red) vs. pooled non-proestrus (blue). Right, mean z-scored activity of selected clusters during indicated social behaviors.

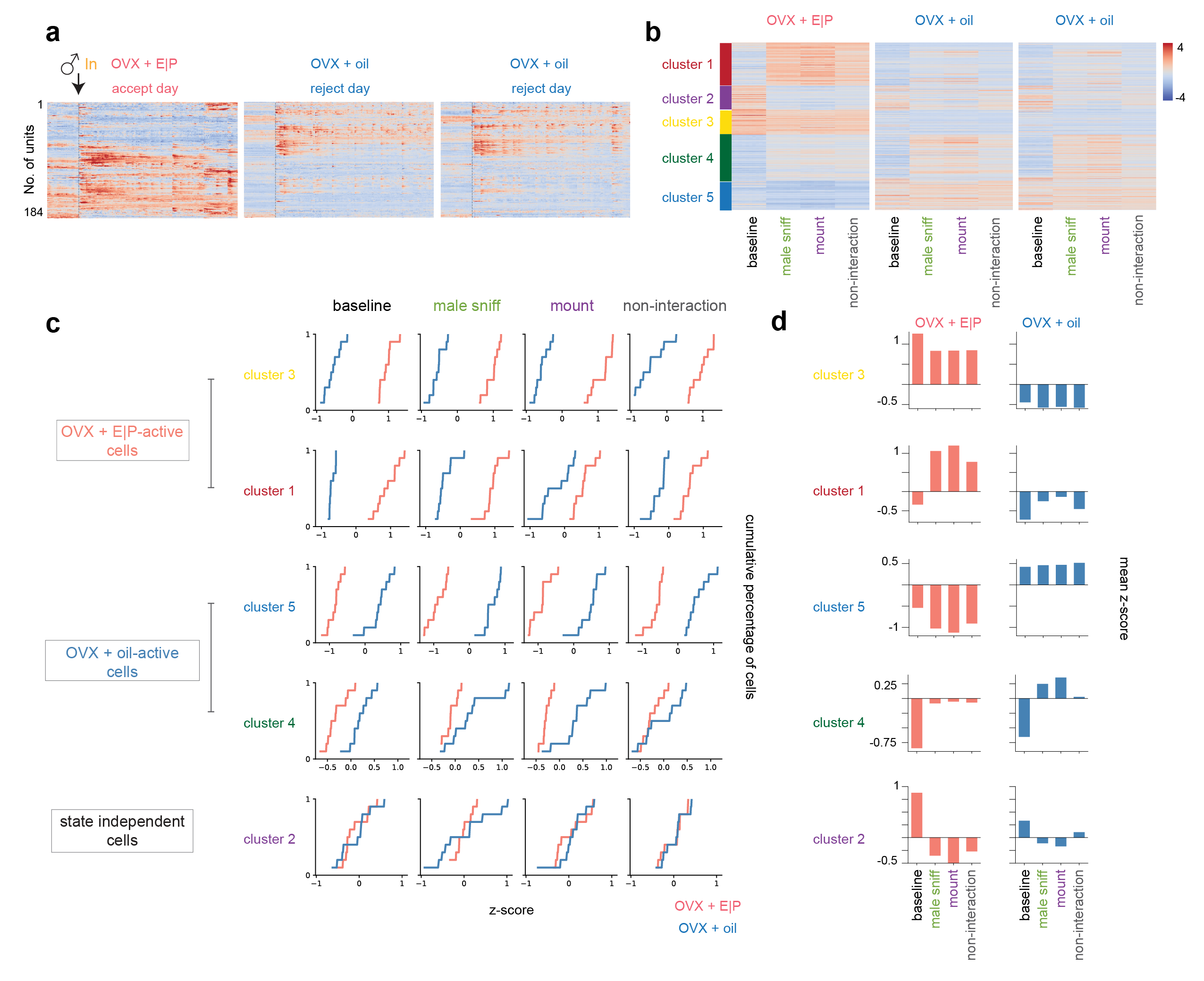
**Extended Data Fig.6 | Single cell analysis for OVX females, additional information for Fig.3. a,** Example neural traces from one ovariectomized longitudinally imaged female. **b,** Unsupervised clustering of neuronal activity in ovariectomized females during interactions with a male in hormone primed (E|P) and unprimed (“oil”) states. Combined data from N = 4 mice. Units z-scored across days. Heat map indicates standard deviations from the mean (computed across all frames). **c,** Cumulative distribution of mean neuronal activity in indicated clusters during different social behaviors. Colors indicate hormone primed (red) vs. unprimed (blue) states. **d,** Mean z-scored activity of selected clusters during indicated social behaviors.

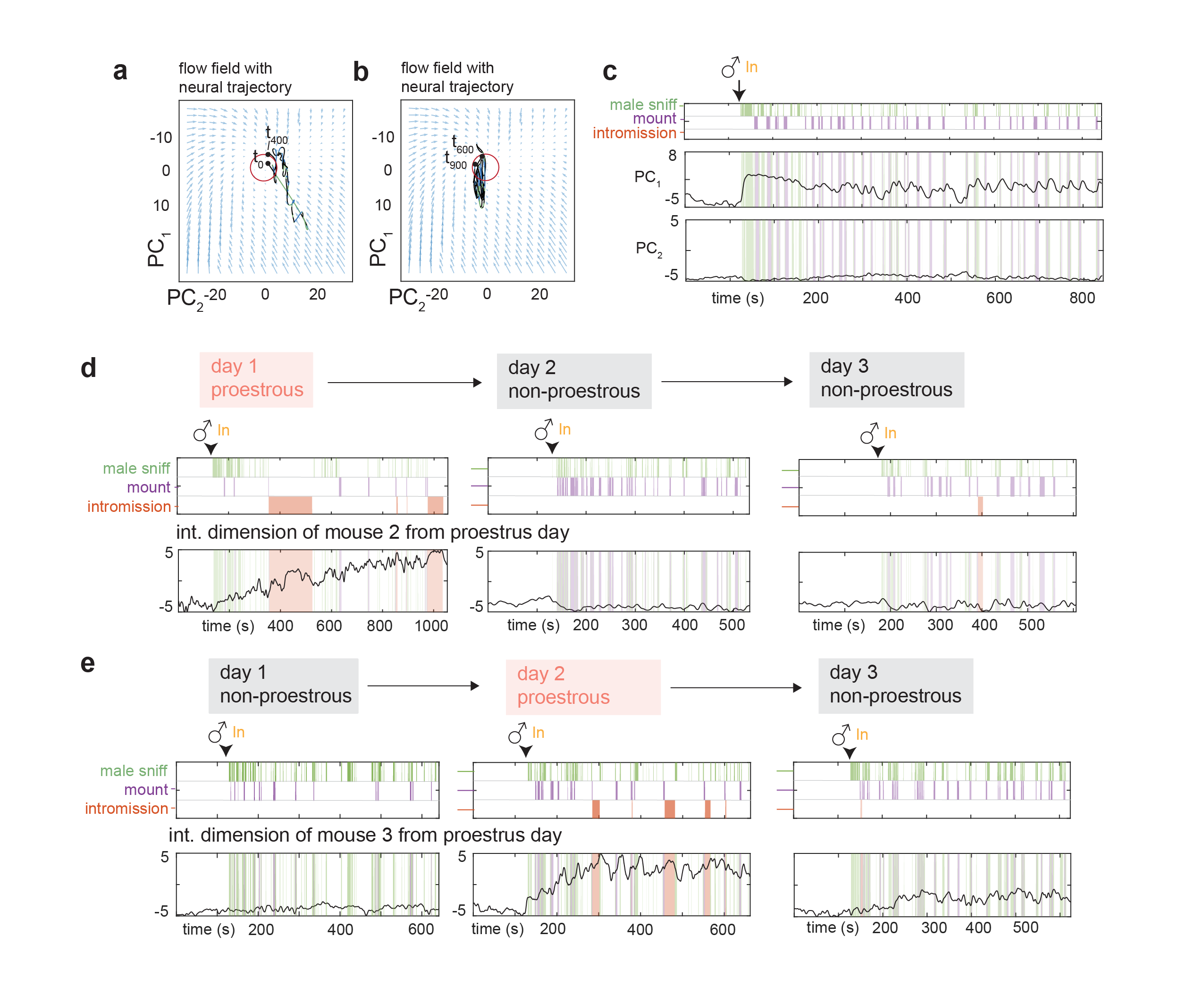
**Extended Data Fig.7 | Dynamical system modelling in non-proestrus days, additional information for Fig.4. a,** Neural trajectories and flow field of VMHvl dynamical system during non-proestrus day for t = 0 to t = 400s. **b,** Same as for t = 400s to t = 900s. **c,** Low dimensional principal components of VMHvl α dynamical system in proestrus day of mouse 1 with neural data projected from non-proestrus day. **d,** Dynamics of integration dimension in VMHvl mouse 2 discovered during proestrus day compared to activity of the same dimension on non-proestrus days. **e,** Same as **d** for mouse 3.

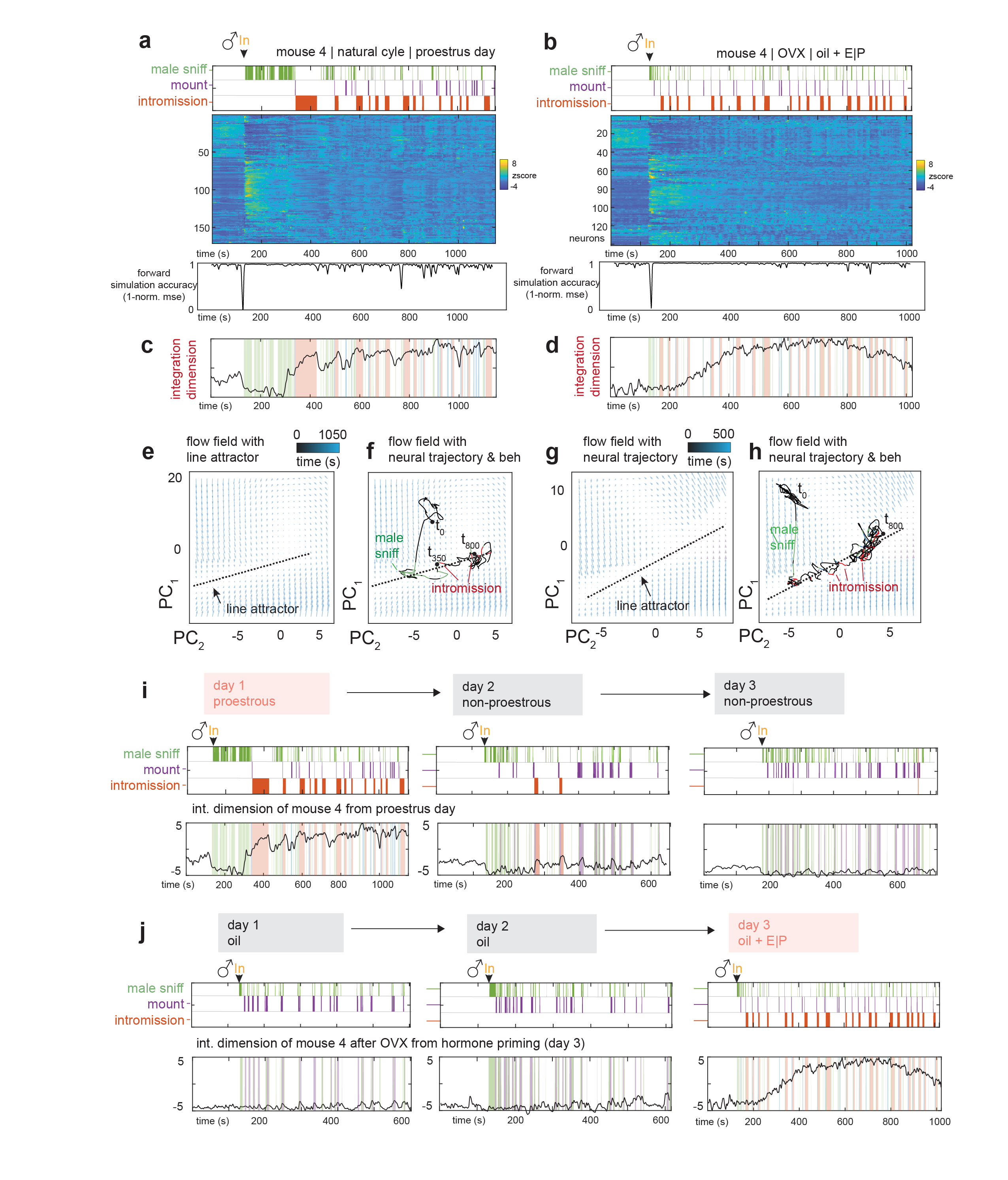
**Extended Data Fig.8 | Natural cycling vs. OVX hormone priming, additional information for Fig.4.** (**a, b,**) Neural raster and behaviors and rSLDS model performance (measured as forward simulation error, see Methods) for mouse 4 in proestrus day of natural estrus cycle **a,** and same mouse 4 on hormone primed day after OVX (day3, oil + E|P) **b.** **(c, d,)** Integration dimension identified by rSLDS on proestrus day in mouse 4 **c,** and during hormone primed day after OVX **d**. (**e, f,**) Flow field **e**, and neural trajectories of dynamical system **f**, with line attractor highlighted of model fit during the proestrus state of the estrus cycle. **(g, h,)** Same as **e, f,** for model fit during hormone primed day after OVX. **i,** Dynamics of integration dimension in mouse 4 discovered during proestrus day compared to activity of the same dimension on non-proestrus days. **j,** Dynamics of integration dimension in mouse 4 discovered during hormone primed day (day 3) compared to the activity of the same dimension during non-primed days.

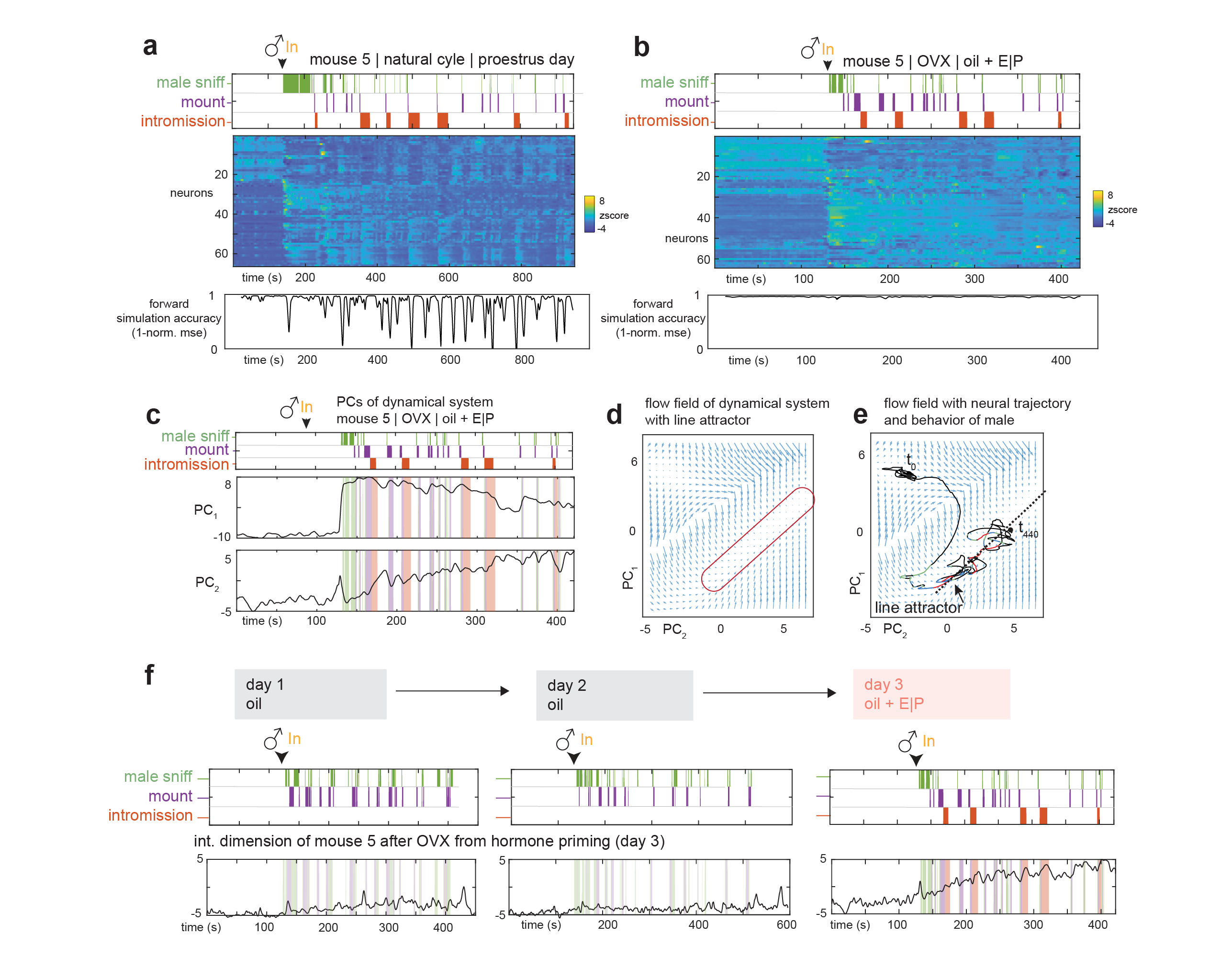
**Extended Data Fig. 9 | Increase of rSLDS model fits after OVX hormone priming.** (**a, b,**) Neural raster and behaviors and rSLDS model performance for mouse 5 in proestrus day of natural estrus cycle **a,** and same mouse 5 on hormone primed day after OVX (day3, oil + E|P) **b**. **c,** Principal components of mouse 5 dynamic system fit during hormone primed day. (**d, e,**) Flow field **d**, and neural trajectories of dynamical system **e**, with line attractor highlighted of model fit during the hormone primed day after OVX in mouse 5. **f,** Dynamics of integration dimension in mouse 5 discovered during hormone primed day (day 3) compared to the activity of the same dimension during non-primed days.

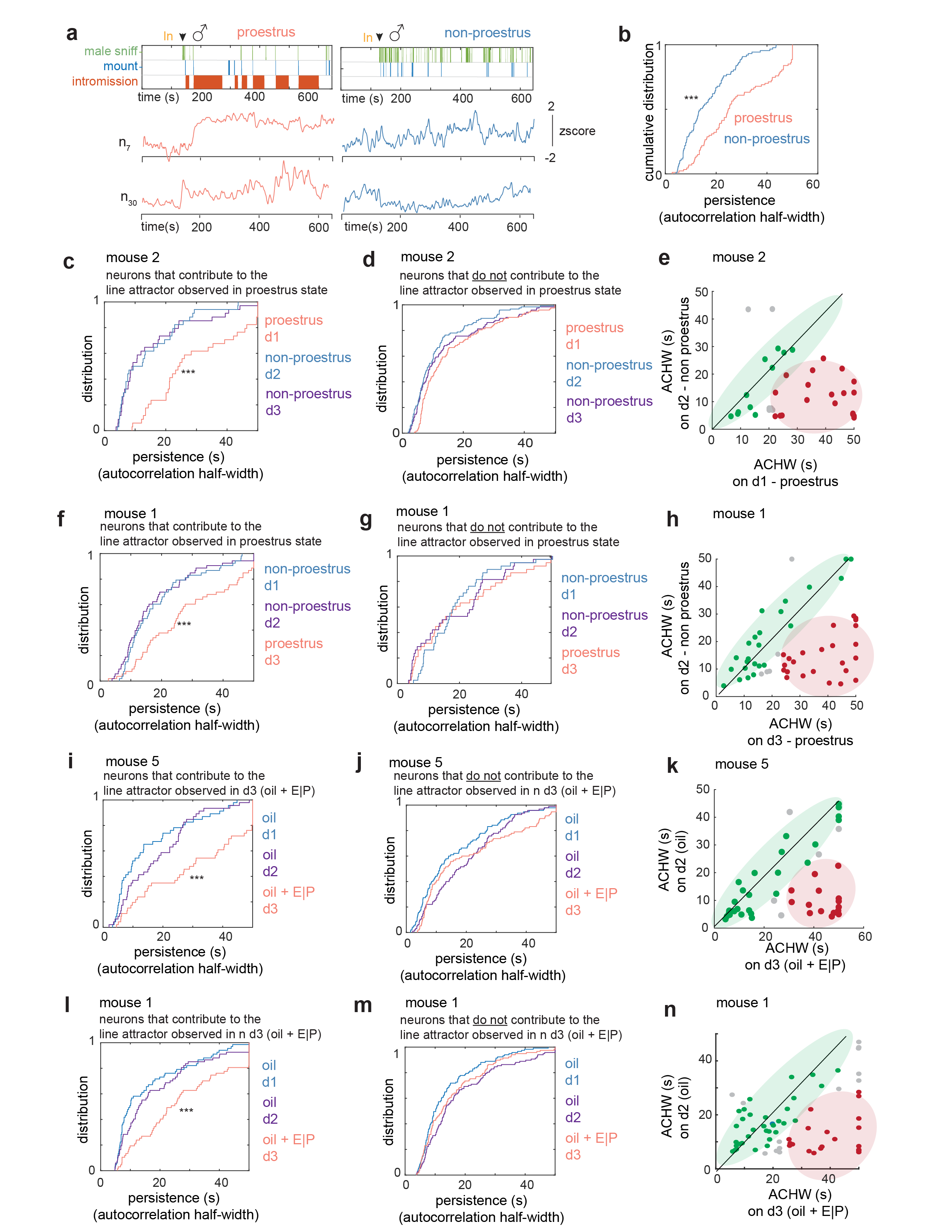
**Extended Data Fig. 10 | Line attractor cells display hormonal state-dependent persistence. a,** Example units active during both proestrus (red traces, left) and non-proestrus (blue traces, right), showing persistence on proestrus day and fast dynamics on the non-proestrus days. **b,** Comparison of cumulative distribution of ACHWs to that of same neurons on non-proestrus days. Data from mouse 1. **c,** Cumulative distribution of ACHWs for units with significant weights on integration dimension across proestrus and non-proestrus day. Data from mouse 1. **d,** Cumulative distribution of ACHWs for mouse 1, for units that do not contribute to the integration dimension on the proestrus day, compared on proestrus vs non-proestrus days. **e,** Scatter plot of ACHWs for units with significant weights on integration dimension for proestrus day vs non-proestrus day. Data from mouse 1. (**f-h**) Same as **c-e** for mouse 2. **i,** Cumulative distribution of ACHWs for units with significant weights on integration dimension across hormone primed (day 3) and non-primed days (days 2, 1). Data from mouse 5. **j,** Cumulative distribution of AHWs for mouse 1, for units that do not contribute to the integration dimension across hormone primed (day 3) and non-primed days (days 2, 1). Data from mouse 5. **k,** Scatter plot of ACHWs for units with significant weights on integration dimension for hormone-primed day vs non-primed day. Data from mouse 1. (**l-n**) Same as **i-j** for mouse 1.
